# Fresh extension of *Vibrio cholerae* competence type IV pili predisposes them for motor-independent retraction

**DOI:** 10.1101/2021.03.09.434644

**Authors:** Jennifer L. Chlebek, Triana N. Dalia, Nicolas Biais, Ankur B. Dalia

**Affiliations:** Department of Biology, Indiana University, Bloomington, IN 47405; Biology Department and Graduate Center, City University of New York, Brooklyn, New York

## Abstract

Bacteria utilize dynamic appendages called type IV pili (T4P) to interact with their environment and mediate a wide variety of functions. Pilus extension is mediated by an extension ATPase motor, commonly called PilB, in all T4P. Pilus retraction, however, can either occur with the aid of an ATPase motor, or in the absence of a retraction motor. While much effort has been devoted to studying motor-dependent retraction, the mechanism and regulation of motor-independent retraction remains poorly characterized. We have previously demonstrated that *Vibrio cholerae* competence T4P undergo motor-independent retraction in the absence of the dedicated retraction ATPases PilT and PilU. Here, we utilize this model system to characterize the factors that influence motor-independent retraction. We find that freshly extended pili frequently undergo motor-independent retraction, but if these pili fail to retract immediately, they remain statically extended on the cell surface. Importantly, we show that these static pili can still undergo motor-dependent retraction via tightly regulated ectopic expression of PilT, suggesting that these T4P are not broken, but simply cannot undergo motor-independent retraction. Through additional genetic and biophysical characterization of pili, we suggest that pilus filaments undergo conformational changes during dynamic extension and retraction. We propose that only some conformations, like those adopted by freshly extended pili, are capable of undergoing motor-independent retraction. Together, these data highlight the versatile mechanisms that regulate T4P dynamic activity and provide additional support for the long-standing hypothesis that motor-independent retraction occurs via spontaneous depolymerization.

**SIGNIFICANCE:** Extracellular pilus fibers are critical to the virulence and persistence of many pathogenic bacteria. A crucial function for most pili is the dynamic ability to extend and retract from the cell surface. Inhibiting this dynamic pilus activity represents an attractive approach for therapeutic interventions, however, a detailed mechanistic understanding of this process is currently lacking. Here, we use the competence pilus of *Vibrio cholerae* to study how pili retract in the absence of dedicated retraction motors. Our results reveal a novel regulatory mechanism of pilus retraction that is an inherent property of the external pilus filament. Thus, understanding the conformational changes that pili adopt under different conditions may be critical for the development of novel therapeutics that aim to target the dynamic activity of these structures.

## INTRODUCTION

Bacteria utilize external hair-like appendages called type IV pili (T4P) to mediate a wide variety of functions including biofilm formation, interbacterial interactions, attachment, horizontal gene transfer, twitching motility, and virulence (1-9). Pili are dynamic structures that extend and retract from the bacterial cell and this dynamic activity is important for many of these functions (1, 2, 10-12). Pili are almost entirely composed of a single repeating subunit called the major pilin protein (13). In Gram-negative bacteria, a complex nanomachine assembles membrane-anchored major pilin subunits into a helical pilus filament that sits atop a platform that is anchored in the inner membrane (14). This pilus filament is then extruded through the outer membrane secretin pore (15). Canonically, extension and retraction of the pilus occurs through the interaction of cytoplasmic hexameric ATPases with the inner membrane platform, where ATP hydrolysis facilitates the coordinated movement of the platform to incorporate pilin subunits into the growing pilus filament (i.e. pilus extension), or remove pilin subunits from the filament which returns them to the inner membrane (i.e. pilus retraction) (16, 17). The complex pilus nanomachine also comprises other structural components which connect the platform to the secretin and are required for proper pilus biogenesis (15, 18, 19).

All T4P systems encode a predicted extension ATPase, commonly called PilB, that facilitates pilus polymerization (20, 21). Canonically, pilus retraction is mediated by a dedicated retraction ATPase (22, 23), however two other modes of pilus retraction have also been described: a bifunctional ATPase that promotes both pilus extension and retraction (24); and pilus retraction that occurs without the aid of any ATPase motors (12, 25, 26). Many T4P systems likely rely on motor-independent retraction, and even many motor-dependent T4P have recently been shown to retract in the absence of their dedicated motors (10, 11, 26). Thus, motor-independent retraction is likely a conserved feature of diverse T4P. While significant advances have been made in our understanding of motor-dependent retraction (21-23, 27-30), a mechanistic understanding of motor-independent retraction and the factors that regulate this process remains lacking.

A long-standing hypothesis is that motor-independent retraction occurs through the spontaneous disassembly of the pilus filament (31). In support of this hypothesis, recent work from our group demonstrates that reducing the stability of pilin-pilin interactions within the pilus filament can enhance motor-independent retraction (25). Despite the fact that motor-independent retraction is, in part, an inherent property of the pilus filament, it can still be regulated by other components of the pilus machine. For example, in the *V. cholerae* toxin-coregulated pilus, motor-independent retraction is triggered by incorporation of a minor pilin into the growing pilus fiber (12). It is possible that other factors may also regulate the stability of pili and/or trigger spontaneous depolymerization.

To determine other factors that may regulate motor-independent retraction, we employed the competence T4P of *Vibrio cholerae* as a model system. The extension ATPase PilB and the retraction ATPases PilT and PilU drive the dynamic assembly and disassembly of competence pili, respectively. Previous work shows that PilB is required for pilus dynamic activity and that a Δ*pilB* mutant does not produce pili (4, 25, 32). This system has recently been used to study motor-independent retraction because a mutant strain lacking both retraction motors (Δ*pilTU*) is capable of motor-independent retraction at a reduced frequency (10, 22, 25). In order to study the dynamics of motor-independent retraction in this system, we employ two main techniques: live imaging of fluorescently labelled pili using a technique (10, 33) in which an amino acid residue of the major pilin, PilA, is replaced by a cysteine (*pilA*^*S56C*^) for subsequent labelling with maleimide conjugated dyes (34); and functional assessment of competence pilus-dependent DNA uptake by performing natural transformation assays (4, 10). Studying the mechanism and regulation of motor-independent retraction in this pilus system may be broadly applicable to the diverse T4P that naturally exhibit motor-independent retraction.

## RESULTS

### Freshly extended pili are more likely to exhibit motor-independent retraction

While observing the dynamic pilus activity in a Δ*pilTU* strain, we observed that pili exist in three possible states: active extension, active retraction, or static surface pili (i.e. extended pili that are not dynamically extending or retracting). The vast majority of pili in the Δ*pilTU* strain are static surface pili, which manifests as a hyperpiliated phenotype (22, 25). By directly examining the rare motor-independent retraction events in Δ*pilTU* cells, we found that the majority were immediately preceded by pilus extension (∼80%), while static surface pili (which are much more abundant compared to actively extending pili) rarely retract and correspondingly only represent ∼20% of retraction events (**Fig. 1A-B**). This suggested that fresh extension predisposes pili for motor-independent retraction.

**Figure 1.**
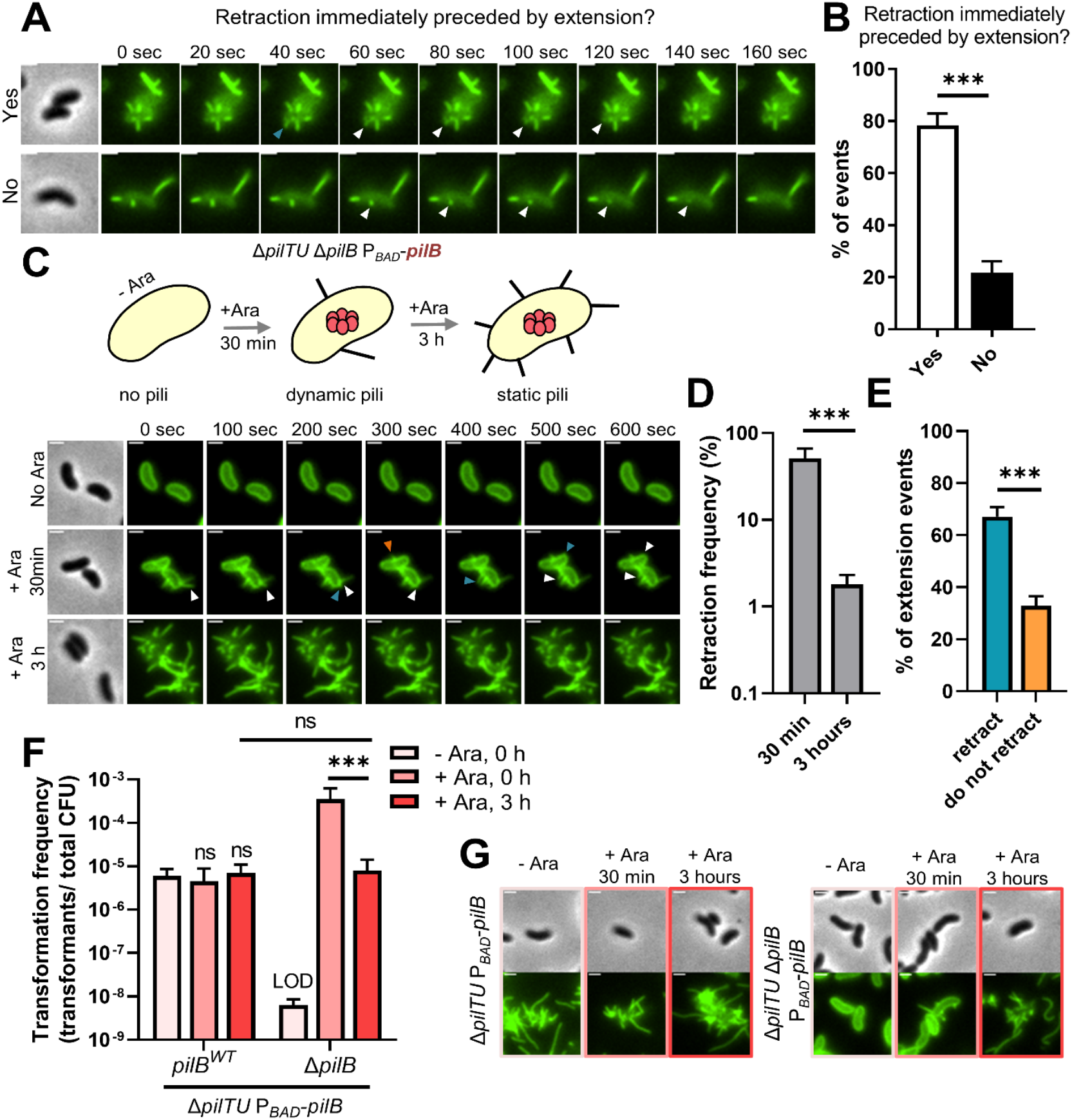
Fresh extension predisposes pili for motor-independent retraction. **(A)** Representative images of Δ*pilTU* retraction events which are either immediately preceded by active extension (Yes) or remain statically extended for ≥30 seconds before retracting (No). An active extension event is indicated with a blue arrow and pilus retraction events are labelled with white arrows. Scale bar, 1 µm. **(B)** Quantification of the percent of Δ*pilTU* retraction events that fall into the categories highlighted in **A**. Data are from 6 independent biological replicates, *n* = 48. **(C)** Schematic (top) of the piliation status of a Δ*pilTU* Δ*pilB* P_*BAD*_-*pilB* strain grown without arabinose (no pili), incubated with 0.2% arabinose for 30 minutes which induces the production PilB shown as a red hexamer (dynamic pili), or incubated with 0.2% arabinose for 3 hours (static pili). Representative montages (bottom) of the dynamic activity of pili at each of these three time points. The blue arrows indicate active pilus extension that is immediately followed by retraction (white arrows). An orange arrow indicates a pilus extension event that does not retract and remains statically extended for the remainder of the time lapse. **(D)** Retraction frequency measured by epifluorescence microscopy of a AF488-mal labeled Δ*pilTU* Δ*pilB* P_*BAD*_-*pilB* strain grown without inducer and then imaged after incubation with 0.2% arabinose for 30 min or 3 hours. Data are presented as the percentage of cells within the population that exhibited a retraction event within the 10 min time-lapse. Data are from three independent biological replicates; 30 min, *n* = 527. 3 hours, *n* = 814. **(E)** Pilus retraction in a Δ*pilTU* Δ*pilB* P_*BAD*_-*pilB* strain incubated with 0.2% arabinose for 30 min was analyzed. Data are presented as the percentage of extension events that ‘retract’ vs ‘do not retract’. Data are from three independent biological replicates; *n* = 263. **(F)** Natural transformation assays of the indicated strains grown without inducer and incubated with or without 0.2% arabinose for the indicated period of time before addition of 200 ng of transforming DNA (tDNA). Both strains for the ‘0 h’ conditions, *n* = 4. ‘3 h’ condition, *n* = 3. **(G)** Representative images of AF488-mal labeled cells to show the piliation status for the specified strains incubated under the conditions indicated. Scale bar, 1 µm. All bar graphs are shown as the mean ± SD. Comparisons in **B, D** and **E** were made by Student’s t-test and comparisons in **F** were made by one-way ANOVA with Tukey’s post test. Ara, arabinose; LOD, limit of detection; ns, not significant; *** = *P* < 0.001.

So next, we sought to test whether increasing the number of pilus extension events could increase the frequency of motor-independent retraction. As mentioned above, a Δ*pilTU* strain is hyperpiliated due to the large number of static surface pili, which may block new extension events from occurring (**Fig. 1A** and **Movies S1-2**) (22, 25). In order to increase the number of extension events, we engineered a Δ*pilTU* Δ*pilB* P_*BAD*_-*pilB* strain where PilB expression can be tightly induced in the presence of arabinose (**Fig. 1C**) (32). This strain was grown without arabinose so that no pili were initially made, and then arabinose was added to induce PilB expression immediately before cells were imaged to track pilus dynamic activity (**Fig. 1C** and **Movies S3-S4**). We also incubated this same strain in the presence of arabinose for 3 hours, which, as expected, became hyperpiliated and phenocopied a Δ*pilTU* mutant (**Fig. 1C** and **Movie S5**) (22). Indeed, we observed many more extension events and the retraction frequency was ∼10-fold greater when PilB was induced immediately prior to imaging compared to when PilB was induced for 3 hours (**Fig. 1C, D** and **Movie S4**). Importantly, the retraction frequency observed after PilB was induced for 3 hours was similar to what has been measured previously for a Δ*pilTU* strain (22, 25). By observing the dynamic events in a Δ*pilTU* Δ*pilB* P_*BAD*_-*pilB* strain after 30-minute incubation with arabinose, we determined that each fresh pilus extension had a ∼65% chance of immediately undergoing motor-independent retraction after extension (**Fig. 1E**). We also wondered if other properties could predict the likelihood that pili would undergo motor-independent retraction like the length of the pilus filament or the piliation status of the cell (i.e. if cells already had one or more extended surface pili). There was no correlation, however, between these features and the likelihood for a pilus to undergo motor-independent retraction (**Fig. S1A, B**). Additionally, cells have a finite number of T4P pilus machines with which to build a pilus fiber (35) and there is a ∼35% chance a pilus will not immediately undergo motor-independent retraction in a Δ*pilTU* mutant and will become statically extended. Eventually, cells will slowly accumulate these static surface pili until there are no T4P machines left to assemble a new pilus, which is consistent with the observed hyperpiliated phenotype (**Fig. 1C, 1G** and **Movie S5**).

These data suggest that the frequency of motor-independent pilus retraction is markedly enhanced when cells extend new pili but that cells will slowly accumulate statically extended pili that fail to retract which will block further extension events. To test this further, we used a transformation assay. We hypothesized that if fresh extension increases the frequency of pilus retraction, then cells that extend more pili should have a higher transformation frequency. To that end, we grew the Δ*pilTU* Δ*pilB* P_*BAD*_-*pilB* strain without arabinose and then added arabinose immediately before transforming DNA (tDNA) was added to reactions (to increase the number of fresh extension events) or grew the same strain and delayed the addition of tDNA until cells were incubated with arabinose for 3 hours (i.e. where cells are already hyperpiliated and therefore exhibiting relatively few fresh extension events) (**Fig. 1C**). We found that the transformation frequency was ∼50-fold higher when PilB was induced immediately before tDNA was added (**Fig. 1F**). As a control, we tested the transformation frequency and piliation status of a Δ*pilTU* P_*BAD*_-*pilB* strain with *pilB* intact and saw that all conditions behaved similarly, indicating that arabinose and/or ectopic expression of PilB does not alter transformation frequency (**Fig. 1F, G)**. Importantly, cells incubated in arabinose for 3 hours showed a similar transformation frequency to the strain where native *pilB* was intact and presented a hyperpiliated phenotype as expected (**Fig 1F, G**). This is consistent with these statically extended pili blocking additional fresh extension events. Altogether, these data show that PilB promotes motor-independent retraction, at least in part, by promoting extension of new pili, which are quantitatively more likely to undergo motor-independent retraction.

### The extension ATPase PilB only enhances motor-independent retraction by promoting fresh pilus extension

Motor-independent retraction seems to be closely coupled with extension and therefore PilB activity. A simple explanation for this would be that PilB is a bifunctional motor powering both extension and retraction in the absence of PilT and PilU (24). However, our recent findings suggest that PilB is not a bifunctional motor because reducing the ATPase activity of PilB did not affect motor-independent retraction rates (25). Although PilB does not power Δ*pilTU* retraction via its ATPase activity, it remains possible that it regulates this process. For example, PilB may halt motor-independent retraction by sitting at the base of the pilus machine and preventing the disassembly of the pilus filament. Conversely, PilB may be required to align the platform to allow for appropriate motor-independent retraction. Therefore, we sought to formally examine whether PilB contributed to motor-independent retraction of *V. cholerae* competence pili. Because PilB is required for pilus extension (4, 25), we cannot test this using a Δ*pilB* mutant. Instead, we sought to specifically degrade PilB after pili were extended. To do this, we adapted a previously described orthogonal degron system derived from the *Mesoplasma florum* tmRNA system (36). Specifically, we C-terminally tagged PilB with a protein degradation tag (pdt2) at the native locus and then ectopically expressed the cognate *M. florum* Lon protease (mf-Lon) using a tightly inducible P_*BAD*_ promoter, which will specifically degrade pdt2 tagged proteins (**Fig. 2A**). Importantly, both *pilB-pdt2* and 3xFLAG-*pilB-pdt2* are both fully functional (**Fig. S2A**). We first assessed the functionality of this degron system by growing *pilB*-*pdt2* P_*BAD*_-*mf-lon* cells consistently in arabinose to induce mf-Lon-dependent degradation of PilB. As expected, strains with *pilB-pdt2* and 3xFLAG-*pilB-pdt2* showed a dramatic reduction in natural transformation specifically when mf-Lon was induced (**Fig. S2A**). Importantly, induction of mf-Lon did not alter the transformation frequency in cells where PilB lacked a pdt2 tag, indicating that mf-Lon does not have a pleiotropic effect on natural transformation (**Fig. S2A**). A western blot also confirmed these results and showed that 3xFLAG-PilB-pdt2, but not 3xFLAG-PilB, is degraded specifically when mf-Lon is expressed (**Fig. S2B**).

**Figure 2.**
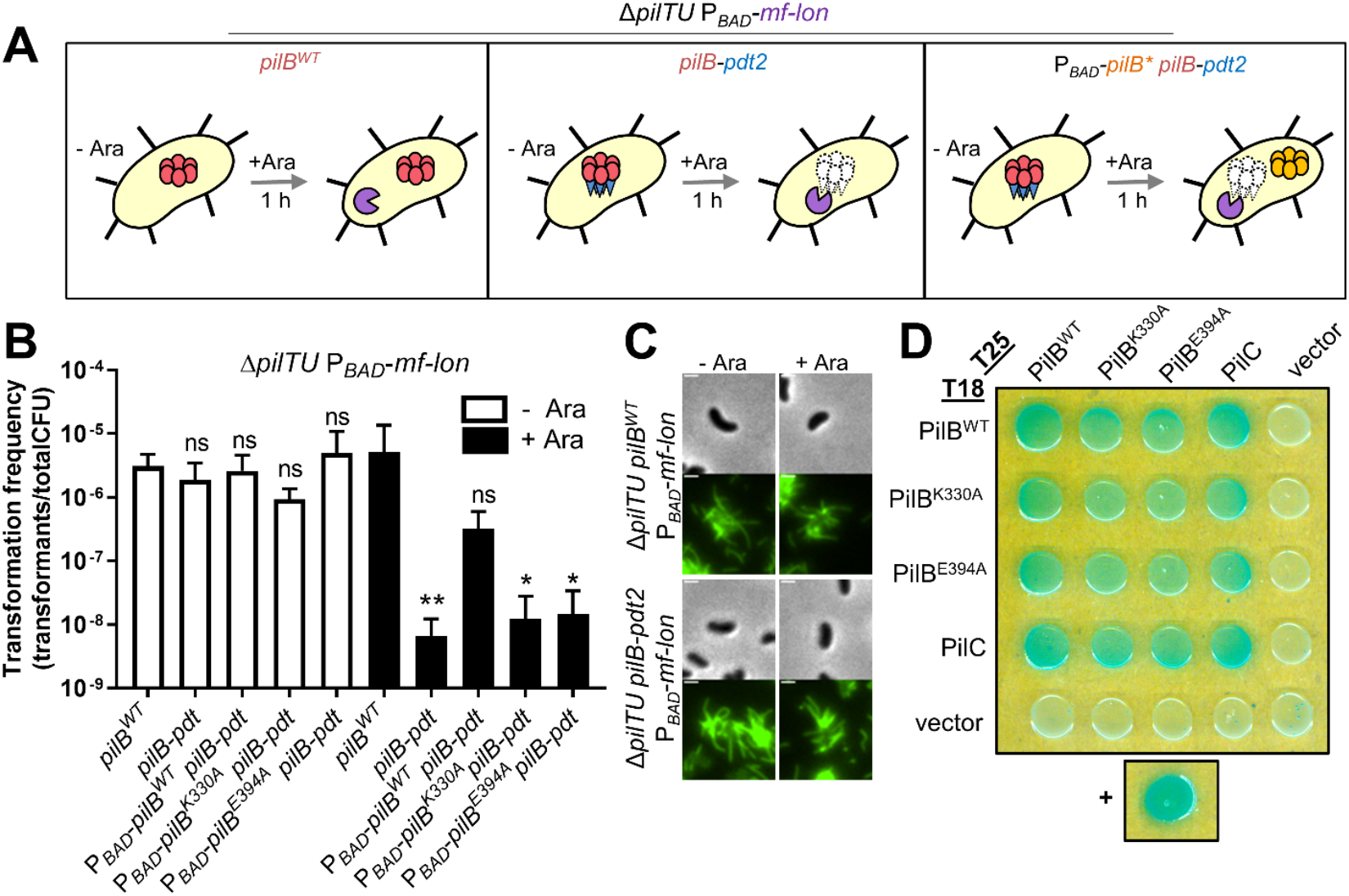
The extension ATPase PilB only enhances motor-independent retraction by promoting fresh pilus extension. **(A)** Schematic of the protein and piliation status of the indicated Δ*pilTU* P_*BAD*_-*mf-lon* strains following incubation with 0.2% arabinose for 1 hour which induces the expression of genes regulated by the P_*BAD*_ promoter. The following proteins are represented in the diagram: mf-Lon protease (purple partial circle), PilB^WT^ (red hexamer), PilB-pdt2 (red hexamer with blue triangles), degraded PilB-pdt2 (white, dashed hexamer and triangles), ectopically-induced untagged *pilB*\* alleles (orange hexamer). Where *pilB*\* denotes different alleles of pilB (*pilB*^*WT*^, *pilB*^*K330A*^, or *pilB*^*E394A*^). **(B)** Natural transformation assays of the indicated Δ*pilTU* P_*BAD*_-*mf-lon* strains grown without inducer and incubated with or without 0.2% arabinose for the 1 h before addition of 200 ng tDNA. All strains, *n* = 3. Bar graphs show the mean ± SD. Asterisk(s) directly above bars denote comparisons to the parent strain. All comparisons were made by one-way ANOVA with Dunnett’s post test. Ara, arabinose; LOD, limit of detection; NS, not significant; * = *P* < 0.05; ** = *P* < 0.01. **(C)** Representative images of the indicated strains incubated with or without 0.2% arabinose for 1 h before imaging. Scale bar, 1 µm. **(D)** Bacterial Adenylate Cyclase Two-Hybrid (BACTH) assay to test pair-wise interactions between T18 and T25-tagged PilB, PilB^K330A^, PilB^E394A^, and PilC. Each construct was also tested for interactions with empty T18 and T25 vectors as a negative control. Leucine zipper fusions to T18 and T25 were used as the positive control (+) and empty T18 and T25 vectors served as an additional negative control. Image is representative of three independent biological replicates.

Using this system, we first grew a Δ*pilTU pilB*-*pdt2* P_*BAD*_-*mf-lon* strain without arabinose to allow for the production of pili, and then subsequently degraded PilB by adding arabinose to induce expression of mf-Lon (**Fig. 2A**). If PilB does not regulate motor-independent retraction, we hypothesized that natural transformation would be unaltered by this treatment, however if PilB contributes to motor-independent retraction we would expect rates of natural transformation to be altered. If PilB inhibits motor-independent retraction, we hypothesized that depletion of PilB should enhance pilus retraction and transformation frequency; while if PilB promotes motor-independent retraction, depletion should decrease the transformation frequency. We found that following degradation of PilB-pdt2, the Δ*pilTU pilB*-*pdt2* P_*BAD*_-*mf-lon* strain was non-transformable (**Fig. 2B**), which indicates that the presence of PilB is required for motor-independent retraction to occur. Importantly, cells were still hyperpiliated following PilB degradation in these experiments, which is consistent with PilB playing an important role in promoting motor-independent retraction (**Fig. 2C**).

The importance of PilB for motor-independent retraction could be due to either 1) PilB extending new pili which are more likely to retract and/or 2) to PilB holding the platform protein PilC in a conformation that promotes pilus depolymerization. We sought to test the later possibility by determining if just the physical presence of PilB is important for motor-independent retraction. To test this, we utilized the PilB degradation system described above in combination with ectopic expression constructs to induce the expression of ATPase inactive alleles of PilB (**Fig. 2A**). The ATPase activity of PilB was inactivated by mutating conserved and critical residues in the Walker A (*pilB*^*K330A*^) or Walker B (*pilB*^*E394A*^) motifs. Importantly, bacterial two-hybrid (BACTH) analysis indicated that both ATPase inactive alleles of PilB still interact with PilC (**Fig. 2D**). If only the presence of PilB was required for motor-independent retraction and not PilB ATPase activity, then we expected that ectopic induction of ATPase inactive PilB would rescue the loss of transformation after degradation of natively expressed PilB-pdt2. As expected, ectopic expression of *pilB*^*WT*^ (P_*BAD*_-*pilB*^*WT*^) rescued the loss of transformation after degradation of PilB-pdt2; however, ectopic induction of P_*BAD*_-*pilB*^*K330A*^ or P_*BAD*_-*pilB*^*E394A*^ did not rescue transformation indicating that the simple binding of PilB to the T4P machine is not sufficient to promote motor-independent retraction (**Fig. 2B**). Additionally, we have recently shown that ATPase-inactive retraction motors cannot hold the pilus platform in a conformation that allows for pilus fiber depolymerization by demonstrating that strains expressing ATPase-inactive alleles of *pilT* or *pilU* (*pilT*^*K136A*^ Δ*pilU* or *pilT*^*K136A*^ *pilU*^*K134A*^) transformed just as poorly as a Δ*pilTU* strain (22). Together, this suggests that inactive motors cannot promote motor-independent retraction by simply aligning the pilus machinery.

The loss of transformation after the degradation of PilB-pdt2 suggests that PilB activity is somehow important for this process, however our data demonstrate that PilB does not power motor-independent retraction (25) and that its presence in the absence of ATPase activity (i.e. ATPase inactive PilB) is not sufficient to drive this retraction. Thus, the primary role that PilB plays to enhance motor-independent retraction may be to promote fresh extension, which predisposes pili to retract as described above. We therefore sought to determine the mechanism by which fresh extension enhances motor-independent retraction.

### Static surface pili are capable of retraction upon induction of dedicated retraction motors

One explanation for why fresh extension may be important for motor-independent retraction could be that non-retracting pili in the hyperpiliated Δ*pilTU* mutant simply represent “broken” pilus machines where the pilus fiber has disengaged from the pilus platform and/or machine in a manner that makes retraction impossible. This may occur in the absence of the retraction motors due to unregulated pilus extension. If this were the case, we hypothesized that these pili should not be capable of retraction even if a retraction motor is ectopically provided. To test this, we generated a tightly inducible ectopic expression construct for PilT (P_*tac*_-riboswitch-*pilT*) that would allow us to delay when the retraction motor was made in a Δ*pilTU* strain (**Fig. 3A**) (32). We also introduced this construct into strains where we could simultaneously degrade PilB (Δ*pilTU pilB*-*pdt2* P_*BAD*_-*mf-lon* P_*tac*_-riboswitch-*pilT*) or not (Δ*pilTU pilB*^*WT*^ P_*BAD*_-*mf-lon* P_*tac*_-riboswitch-*pilT*) to see if the presence of PilB altered the effect of PilT on pilus retraction (**Fig. 3A**). Cells were initially grown without any inducers, then mf-Lon was induced if applicable (to degrade PilB-pdt2), followed by induction of PilT to assess if pili were capable of retraction (**Fig. 3A**). When we performed these experiments, we saw that the vast majority of extended pili were still capable of retraction—whether PilB was present or not (**Fig. 3B** and **Movies S6-S8**). This suggests that the pili that remain extended outside of the cell in a Δ*pilTU* strain are not simply broken or misaligned with the pilus machine and that whatever is preventing these pili from retracting can be overcome by a retraction ATPase. Therefore, there must be something unique about freshly extended pili that predisposes them to retract. Also, because pili still retract following depletion of PilB, this suggests that PilB is not required for motor-dependent retraction by PilT.

**Figure 3.**
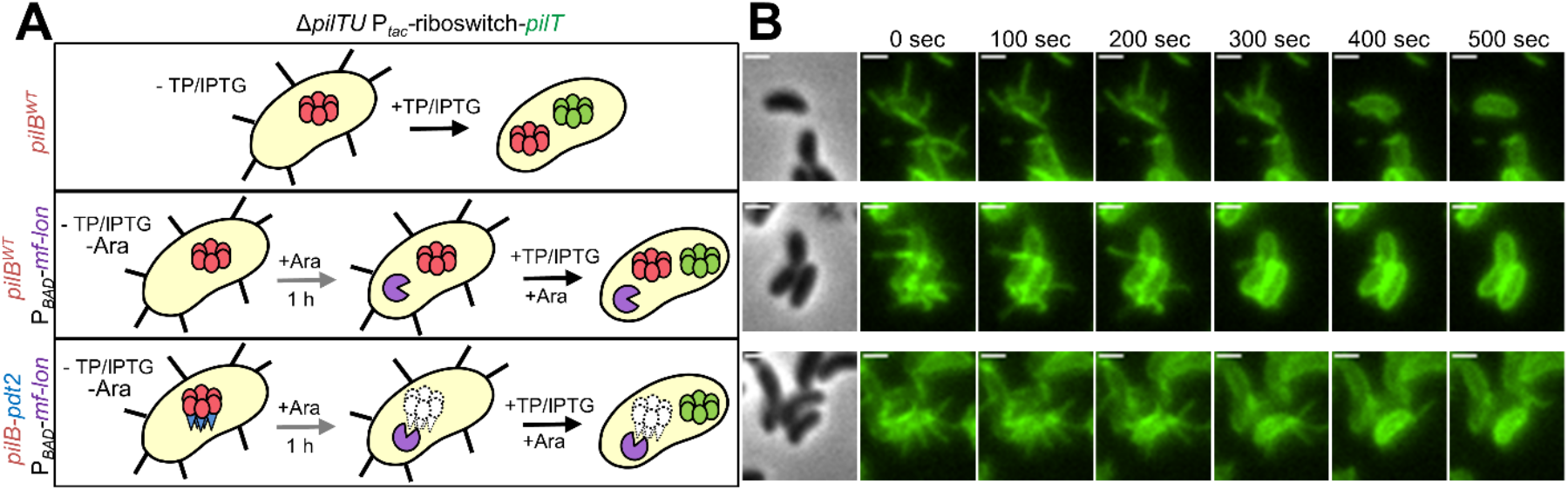
Static surface pili are capable of retraction upon induction of the dedicated retraction motor PilT. **(A)** Schematic of the protein and piliation status of the indicated Δ*pilTU* P_*tac*_-riboswitch-*pilT* strains following growth and incubation conditions with the indicated inducers. Incubation with TP and IPTG induces the P_*tac*_-riboswitch-*pilT* construct. The following proteins are represented in the diagram: PilB^WT^ (red hexamer), ectopically-induced PilT (green hexamer), mf-Lon protease (purple partial circle), PilB-pdt2 (red hexamer with blue triangles), degraded PilB-pdt2 (white, dashed hexamer and triangles). **(B)** Representative montages of the strains / conditions schematized in **A**. All strains were grown without inducers present (-IPTG/TP/Ara) and incubated with the inducers as indicated. The black arrow **A** indicates when imaging started (as depicted in **B**) and the inducers present during the imaging conditions. Scale bar, 1 µm. Ara, arabinose; TP, theophylline; IPTG, Isopropyl β-d-1-thiogalactopyranoside.

### Fresh extension events and fiber instability act through the same pathway to enhance motor-independent pilus retraction

One potential explanation for why pili fail to retract could be that pilus binding to a surface over time prevents retraction due to the relatively smaller retraction forces that motor-independent retraction generates (22). However, cells fail to retract their pili even in liquid culture settings, where cells are not encountering a surface. This is phenotypically observed as hyperpiliation/cell aggregation and a reduced frequency of natural transformation, in liquid culture based assays (22, 25). Thus, attachment of pili to surfaces likely does not influence the ability of freshly extended pili to retract.

Our previous work has revealed that motor-independent retraction is regulated by an inherent property of the pilus fiber in both naturally motor-dependent and motor-independent pilus systems (25). In a suppressor screen aimed at identifying mutations that enhance motor-independent retraction of the *V. cholerae* competence pilus, the only hits identified were in the major pilin gene *pilA*. We show that these *pilA* point mutations (*pilA*^*G34R*^, *pilA*^*V74A*^, *pilA*^*G78S*^ and *pilA*^*G80S*^) likely disrupt pilin-pilin interactions within the pilus filament, which enhanced the frequency of motor-independent retraction (25). Since fresh extension and these *pilA* point mutants both enhance the frequency of motor-independent retraction, we next sought to address whether they acted through the same pathway. To test this, we first used microscopy to measure the percent of pilus retraction events that were immediately preceded by extension in the Δ*pilTU pilA* point mutant backgrounds. We observed that there was a significant increase in the number of retraction events that were not immediately preceded by pilus extension in the Δ*pilTU pilA* point mutants (*pilA*^*G34R*^, *pilA*^*V74A*^, *pilA*^*G78S*^ and *pilA*^*G80S*^) compared to Δ*pilTU pilA*^parent^ (**Fig. 4A**), which suggests that these *pilA* point mutations relieve the need for fresh pilus extension to promote motor-independent retraction. Above, we show that increasing the number of pilus extension events in a Δ*pilTU* strain (via tightly regulated ectopic expression of PilB) increases the transformation frequency (**Fig. 1F**) because freshly extended pili are more likely to retract (**Fig. 1D**). If the *pilA* suppressor mutations truly relieve the need for fresh pilus extension to promote motor-independent retraction, then we hypothesized that they should be epistatic (i.e. that the combination of these features should not yield an additive effect on motor-independent retraction when assessed by natural transformation). Indeed, we observed that inducing fresh extension events in the Δ*pilTU* Δ*pilB* P_*BAD*_*-pilB pilA* point mutant backgrounds immediately before tDNA was added to the transformation assay did not enhance transformation frequency compared to when tDNA was added to these strains after incubation with PilB inducer for 3 hours (**Fig. 4B**). Importantly, the transformation frequency observed in the *pilA* point mutant backgrounds is not the maximum transformation frequency that can be achieved in this assay (**Fig. S2**). Also, induction of fresh extension still enhanced transformation in the Δ*pilTU* Δ*pilB* P_*BAD*_*-pilB pilA*^*parent*^ as previously observed (**Fig. 1F**). Because fresh extension does not enhance retraction in the Δ*pilTU pilA* point mutants, this suggests that fresh extension and the *pilA* point mutants are epistatic and may act via the same mechanism to enhance motor-independent pilus retraction.

**Figure 4.**
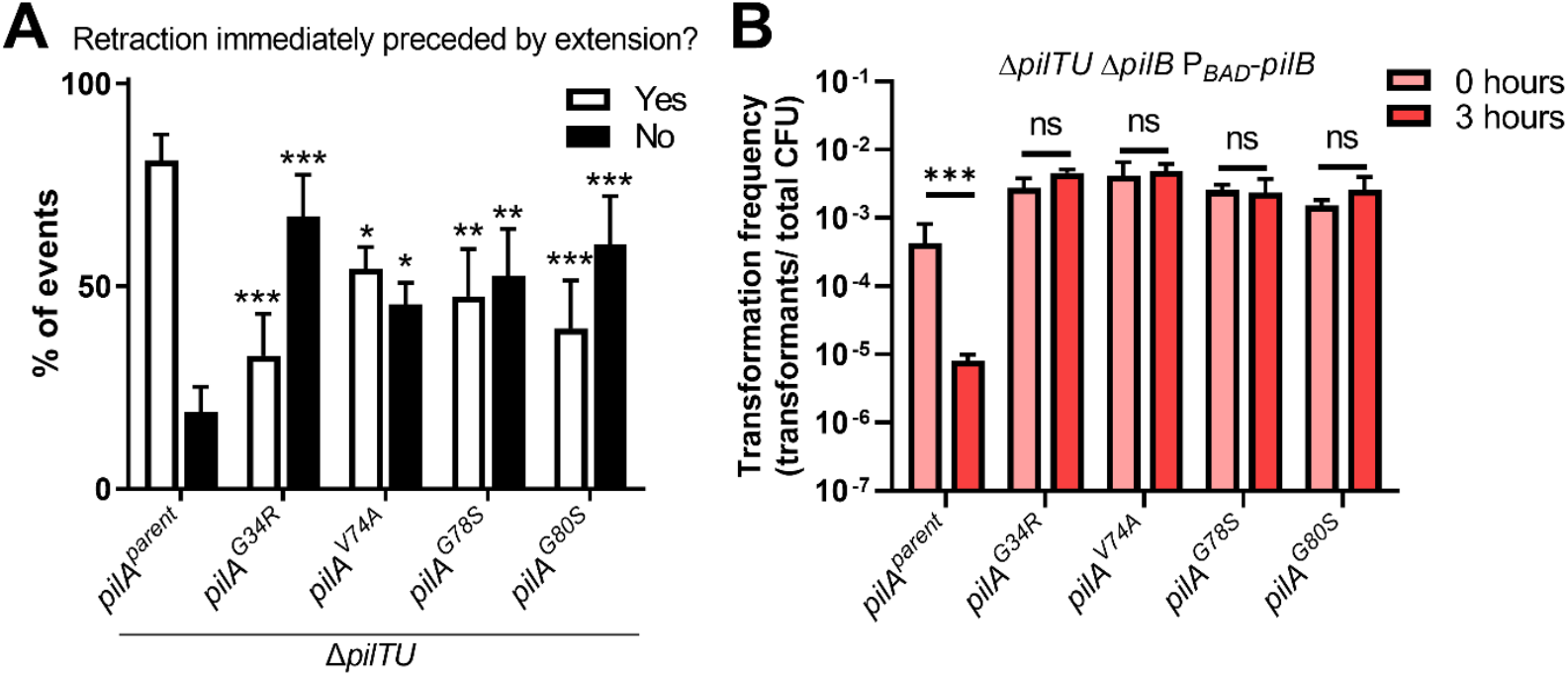
Fresh pilus extension and destabilization of pilus fibers by *pilA* point mutants act via the same pathway to promote motor-independent retraction. **(A)** Quantification of the percent of retraction events in the indicated strains that were either immediately preceded by an extension event (‘Yes’) or occurred in pili that had been statically extended for greater than 30 seconds (‘No’). The *pilA*^*parent*^ data are the same as in **Fig. 1B** and were included here for ease of comparison. Data are from three independent biological replicates: *pilA*^*G34R*^, *n* = 111; *pilA*^*V74A*^, *n* = 246; *pilA*^*G78S*^, *n* = 190; *pilA*^*G80S*^, *n* = 197. **(B)** Natural transformation assays of the indicated P_*BAD*_-*pilB* Δ*pilB* Δ*pilTU* strains grown without inducer and incubated with 0.2% arabinose for the specified period of time before addition of 200 ng of transforming DNA (tDNA). All strains, *n* = 3. All bar graphs are shown as the mean ± SD. Asterisks directly above bars denote comparisons to the parent strain. All comparisons were made by one-way ANOVA with Tukey’s post test. ns, not significant; *** = *P* < 0.001.

### Pilus fibers can adopt unique conformations and may transition between unstable and stable states during and after pilus extension

Our previous work suggests the *pilA* point mutations enhance the frequency of pilus retraction by destabilizing the pilus filament and that fiber instability is a requirement for spontaneous depolymerization during motor-independent retraction (25). Since the effect of fresh extension and the *pilA* point mutations are epistatic, we hypothesized that the pilus filament extends in a less stable state. If the pilus does not immediately retract, it can transition to a more stable conformation that cannot undergo motor-independent retraction (i.e. generating a static surface pilus). For this to be true, the interaction between pilin subunits within the pilus fiber must be able to change to transition the fiber between these distinct unstable and stable states. It has previously been demonstrated that pilus fibers can exist in different structural forms (37-39). Specifically, T4P from *Neisseria gonorrhoeae* can undergo force-dependent structural transitions where the arrangement of pilin subunits within the fiber take on distinct, yet reversible, conformations (40). We hypothesize that freshly extended competence pili in *V. cholerae* may also adopt a distinct conformation that is more permissive for motor-independent retraction.

To start, we first investigated whether pilin subunits within the competence pilus are capable of altering their arrangement within the pilus fiber. One hallmark that would indicate that pilins are capable of undergoing this type of transition, is that the fiber itself should exhibit elastic properties, which would be observed as stretching or lengthening of the pilus fiber under force (40). If pilins were incapable of being easily arranged, then we would expect the filament to be inelastic. To test this, purified fluorescently-labelled pili were tethered between a magnetic bead and an elastic pillar and length of pili was determined by imaging pili by fluorescence microscopy (**Fig. 5A**,**B**). Force was applied to the pilus by turning on a magnet to pull on the tethered bead as previously described (40). When the magnet was “ON” to pull on the pilus fibers, we observed that the length of the pilus increased significantly (∼1.5-fold) compared to pilus fiber when the magnet was turned “OFF” (**Fig. 5C)**. These data show that the competence pilus fiber is elastic and can adopt more than one structural configuration because pilin-pilin interactions within a stretched fiber must be distinct from unstretched fibers to accommodate the increased length. This observation is consistent with our hypothesis that pili may adopt distinct conformations. It is uncertain, however, if pili can undergo these rearrangements during active extension and/or retraction *in vivo*.

**Figure 5.**
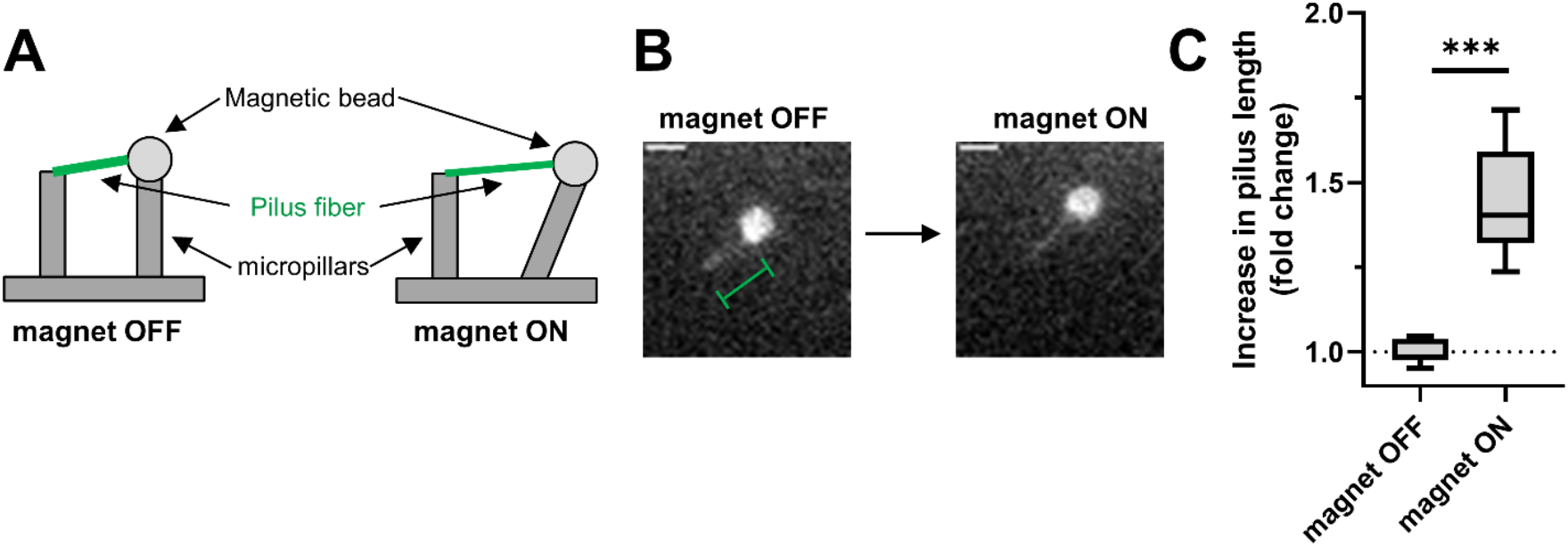
Pilin subunits can undergo rearrangements within the filament as indicated by stretching of pilus fibers. **(A)** Schematic of the experimental design showing a pilus fiber (green) tethered to a magnetic bread (light grey) and a micropillar (dark grey). When a magnet is turned on the magnetic bead is pulled in one direction which stretches the pilus fiber. **(B)** Representative fluorescent images of a fluorescently labeled pilus fiber before stretching (magnet OFF) and after stretching (magnet ON). The magnetic bead can be seen as a white circle in the image and the fluorescent pilus length was measured as highlighted by the green brackets. Scale bar, 1 μm. (C) Quantification of the increase in pilus length relative to ‘magnet OFF’ condition (no force applied). Data are from three independent experiments. Box plots represent the median and the upper and lower quartile, while the whiskers demarcate the range. Comparisons were made by Student’s T-test. *** = *P* < 0.001.

## DISCUSSION

This study sheds light on the mechanisms that regulate motor-independent retraction of the competence T4P of *V. cholerae*. Our findings indicate that newly extended pili are more likely to undergo motor-independent retraction. Based on our data, we propose a model in which the pilus fiber extends in a transiently unstable state which allows for spontaneous depolymerization immediately after extension is complete (**Fig. 6**). However, if a pilus does not retract immediately, pilins within the fiber may reposition into a more stable conformation, which blocks the extended pilus from undergoing motor-independent retraction (**Fig. 6**, *static pilus*). This stable state may be more thermodynamically favorable which further prevents the transition back to a retractable unstable state. Alternatively, rearrangements of pilin-pilin interactions may not be occurring along the entire length of the pilus fiber. The stability of the interaction between the terminal pilin and the pilins at the base of the pilus fiber may be the most important determinant for retraction by spontaneous depolymerization (25, 31). If this is the case, then our data indicate that freshly extending pili have a ∼65% chance of terminating extension in a state where the terminal pilin is poorly interacting with the base of the filament, which would allow for spontaneously depolymerization. While ∼35% of the time, the terminal pilin is stably associated with the base of the pilus fiber, which prevents spontaneous depolymerization and generates a static surface pilus. Ideally, we would like to test the stability and/or arrangement of pilins in freshly extended filaments to static surface pili. Unfortunately, the transient nature of the freshly extended state, and the subtle conformational changes that are likely associated with these phenotypes makes this impossible to test using currently available approaches.

**Figure 6.**
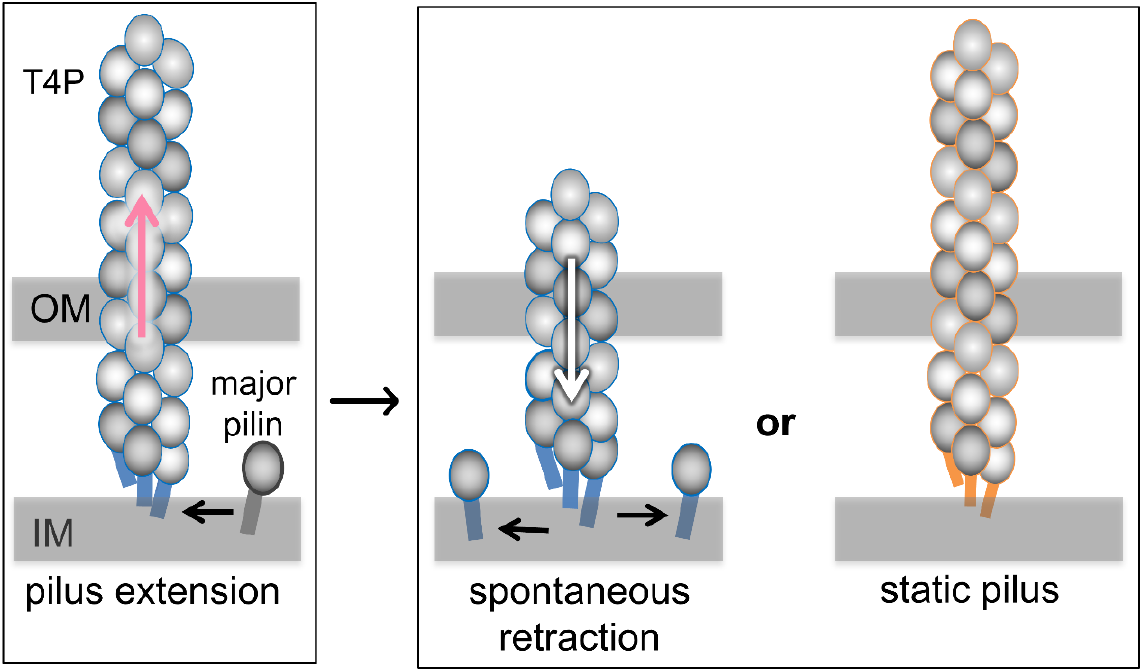
Model for how fresh pilus extension may enhance motor-independent retraction of T4P. In a motor-independent retraction system, the entire pilus fiber may be extended in a less stable state (blue fiber) which allows for spontaneous depolymerization (white arrow) to occur immediately following extension (about ∼65% of the time for the *V. cholerae* competence pilus). If the pilus relaxes into a more stable state (orange fiber) before spontaneous depolymerization occurs, however, this will result in pili remaining statically extended outside the cell. Our data indicates that these static surface pili are not permissive for spontaneous depolymerization (orange fiber) but can be retracted by the PilT motor. An alternative model is that the terminal pilin (black) is simply added to the pilus fiber in an unstable state which allows for spontaneous depolymerization (white arrow); conversely, if the terminal pilin is added to the base of the pilus in a more stable conformation, this may prevent motor-independent retraction. OM, outer membrane; IM, inner membrane.

Aside from our observations that competence T4P can undergo pilin rearrangement within the pilus fiber *in vitro* (**Fig. 5**), strong evidence to support our model (**Fig. 6**) remains lacking. This model is largely bolstered by a previous suppressor screen where the only mutations that could enhance motor-independent retraction all mapped to the *pilA* gene (25). These mutations disrupted the stability of the pilus fiber to enhance motor-independent retraction, which suggested that the mechanism underlying retraction was an inherent property of the pilus fiber. Because we found that fresh pilus extension and these *pilA* point mutations are epistatic, we hypothesize that freshly extended pili must adopt a distinct conformation that is permissive to retraction. The *pilA* point mutations may simply extend the duration of time pili spend in the unstable state following extension. Alternatively, they may alter pilin-pilin interactions within the fiber in a way that entirely prevents the transition into a stable retraction-nonpermissive state. Regardless, these data provide more evidence that the mechanism of motor-independent retraction is an inherent property of the pilus fiber which strongly supports the model that this process occurs via spontaneous depolymerization of the pilus fiber.

So far, our data show that properties that are inherent to the pilus fiber are important for regulating motor independent retraction. Although our data demonstrate that PilB is not required for motor-independent retraction via its ATPase activity (25) or via its presence to align the T4P machine (**Fig. 2**), other T4P components may play an important role in regulating this spontaneous depolymerization (e.g. PilM, PilN, PilO, PilP, and PilC). Future work will be needed to determine if these other factors regulate motor-independent retraction as well.

Our results provide indirect evidence to suggest that pilus filaments adopt distinct structural conformations during active extension and/or retraction. These distinct structural conformations of pilus filaments may be important to the function of pilus systems. Specifically, we show that fresh extension is important for motor-independent retraction in the competence pilus of *V. cholerae*. Although these results were demonstrated in an artificial system (since the *V. cholerae* competence T4P naturally relies on PilTU motors for retraction), recent work reveals that even WT *V. cholerae* harbors many static surface competence pili in its native environmental niche, a chitin biofilm (5). One possible explanation for this phenomenon is that the activity of the PilTU motors are attenuated under these chitin biofilm conditions. Thus, leaving spontaneous depolymerization as the only mechanism for pilus retraction within a chitin biofilm. Furthermore, another recent study suggests that T4P-mediated bacterial movement inside a biofilm is important for biofilm fluidity and bacterial survival (41). With these findings in mind, the motor-independent retraction of *V. cholerae* competence pili following fresh pilus extension observed here may be an important failsafe mechanism for maintaining pilus retraction to support transformation and proper cellular fluidity within a chitin biofilm. Additionally, other examples of pilus systems, such as the toxin co-regulated pilus of *V. cholerae* (42), which naturally exhibits motor-independent retraction, and the T4P of *N. gonorrhoeae* (43) and *Thermus thermophilus* (44), which naturally exhibit motor-dependent retraction, have all been shown to harbor many static surface pili, similar to competence T4P in a Δ*pilTU V. cholerae* strain (**Fig. 1A**). If T4P filaments of other systems also exhibit distinct conformational states in actively extending pili vs static surface pili, it is possible that these states are important for the diverse functions carried out by these pili. It remains unclear if the findings described here are broadly conserved properties of other T4P and testing this hypothesis will be the focus of future work.

Interestingly, the retraction force previously measured for PilT-dependent retraction of the *V. cholerae* competence pilus is ∼10 pN (10, 22) which is the same force used to stretch pili *in vitro* (40) (**Fig. 5C**). This suggests that a retraction motor may be able to ‘pull’ on the pilus with enough force to transition the fiber from a nonpermissive conformation into a retraction permissive conformation. Indeed, our data demonstrate that while static surface pili cannot undergo motor-independent retraction, they can still be retracted by the PilT motor (**Fig. 3** and **Movies S6-S8**). Furthermore, using an orthogonal degron system and our recently characterized tightly inducible ectopic expression constructs (32, 36), we observed that PilT is capable of retracting statically extended pili and that this activity is not affected by the presence or absence of PilB (**Fig. 3** and **Movies S6-S8**). To our knowledge this is the first evidence to suggest that retraction motors can mediate pilus depolymerization in the absence of their cognate extension motors. It was previously shown by many groups that PilB does not require the presence of PilT to mediate pilus polymerization (because retraction motor mutants accumulate many surface pili) (11, 26, 45). However, determining whether the retraction motor requires the presence of the extension motor PilB could not be assessed without the tools established in this study. By allowing for the tightly regulated depletion and expression of pilus components, these tools should prove broadly valuable for addressing fundamental questions about T4P biogenesis and regulation.

## ACKNOWLEDGEMENTS

We would like to thank Lisa Craig for helpful discussions, Julia C van Kessel for providing strains for BACTH assays, Kim Seed for generously providing DNA with the P_*tac*_- riboswitch construct, and James J. Collins for providing plasmids encoding pdt2 and mf-Lon. This work was supported by Grant R35GM128674 from the National Institutes of Health (to ABD) and Grant SC2AI116566 from the National Institutes of Health (to NB).

## MATERIALS AND METHODS

### Bacterial strains and culture conditions

*Vibrio cholerae* and *Escherichia coli* strains were routinely grown in LB Miller broth and on LB Miller agar supplemented with erythromycin (10 μg/mL), kanamycin (50 μg/mL) and spectinomycin (200 μg/mL), carbenicillin (100 μg/mL or 20 μg/mL), trimethoprim (10 μg/mL), chloramphenicol (2 μg/mL), and zeocin (100 μg/mL) when appropriate.

### Construction of mutant strains

*V. cholerae* strains used throughout this study are derivatives of the El Tor isolate E7946. All mutant constructs were generated by splicing-by-overlap extension (SOE) PCR and then introduced into strains via MuGENT and natural transformation exactly as previously described (46-49). The *araC*–P_*BAD*_ region for our P_*BAD*_ constructs was amplified from pBAD18-Kan (50). The P_*tac*_- riboswitch construct was amplified from DNA generously provided by Kim Seed (51). For a detailed list of all mutant strains used throughout this study see **Table S1**. For a detailed list of all primers used to construct mutant strains see **Table S2**.

### Natural transformation assays

In order to induce maximal competence in *V. cholerae* strains, the master competence regulator TfoX is overexpressed using an IPTG-inducible P_*tac*_ promoter and the cells are genetically locked in a regulatory state that mimics high cell density via deletion of *luxO*, as previously described (47, 52-56). Chitin-independent transformation assays were performed exactly as previously described (48). Briefly, strains were grown overnight with aeration at 30 °C then grown to late log phase by subculturing ∼10^8^ colony forming units (CFU) into 3 mL of LB supplemented with 100 µM IPTG, 20 mM MgCl_2_, 10 mM CaCl_2_, +/- 0.2% arabinose as indicated. Next, ∼10^7^ CFU of this culture was diluted into Instant Ocean medium (IO) (7 g/L; Aquarium Systems) supplemented with 100 µM IPTG and 0.2% arabinose was added as indicated. Then, 200 ng of transforming DNA (tDNA) was added to each reaction either immediately or after 1-3 hours of static incubation where indicated. All reactions were then incubated statically at 30 _o_C overnight. The tDNA replaces VC1807, a frame-shifted transposase, with an erythromycin resistance antibiotic marker (aka ΔVC1807::Erm^R^) as previously described (46). Negative control reactions where no tDNA was added were performed for each strain. After incubation with tDNA, reactions were outgrown by adding 1mL of LB to each reaction and shaking (250 rpm) at 37 °C for ∼2 hours. Reactions were then plated for quantitative culture onto agar plates selecting for transformants (LB + 10 μg/mL erythromycin) or onto plain LB for total viable counts. The transformation frequency is defined as the number of transformants divided by the total viable counts. For reactions where no transformants were obtained, a limit of detection was calculated and plotted.

### Pilin labelling, imaging and quantification

In order to label *V. cholerae* competence pili for observation by epifluorescence microscopy, strains were grown to late log exactly as described above for natural transformation assays. Then, ∼10^8^ CFU were spun down at 18,000 x g for 1 min and resuspended in IO. Cells were then incubated with 25 μg/mL AF488-mal for 15 minutes statically at room temperature in the dark. Cells were washed twice, fully disrupting the pellet using 100 μL of IO for each wash, and finally resuspended in IO. Where indicated, 0.2% arabinose was added to cells resuspended in IO and incubated for either 30 min, 1 hour, or 3 hours before imaging. For induction of P_*tac*_-riboswitch-*pilT*, 1.5 mM theophylline and 1 mM IPTG was added to cells immediately prior to imaging.

For imaging, 2μL of labeled cells were placed under an 0.2% gelzan pad on a coverslip and imaged. For the induction of P_*BAD*_- promoters, the gelzan pad was made in IO + 0.2% arabinose where indicated. For the induction of P_*tac*_-riboswitch-*pilT*, the gelzan pad was made in M9 minimal medium (1X M9 salts (Difco), 2 mM MgSO_4_, 0.1 mM CaCl_2_, and 50 μM FeSO_4_) supplemented with 1% glucose, 1.5 mM theophylline, 1 mM IPTG, +/- 0.2% arabinose where indicated. All imaging was done on an inverted Nikon Ti-2 microscope with a Plan Apo ×60 objective, a green fluorescent protein filter cube, a Hamamatsu ORCAFlash 4.0 camera and Nikon NIS Elements imaging software. Representative images of the piliation state of strains were gathered and the lookup tables for each phase or fluorescent image were adjusted to the same range in each figure.

In order to measure dynamic events, labelled cells were imaged by time-lapse microscopy. A phase-contrast image (to image cell bodies) and fluorescent image (to image labeled pili) were taken every 10 seconds for 10 minutes. Time-lapses from 3 independent biological replicates were sectioned into areas containing ∼100-300 cells. In order to calculate the frequency of retraction, the number of cells that exhibited a pilus retraction event and the total number of cells in the field were manually counted. In order to calculate the percent of extension events that retract, the total number of extension events were manually counted and then retraction was manually assessed. Pili were denoted as ‘do not retract’ if no retraction was observed in the given time frame. Pili that finished extending ≤2 frames before the time-lapse concluded were not analyzed. Representative images were prepared for figures using Fiji v2.0.0 (57) and NIS Elements analysis software.

### Bacterial Adenylate Cyclase Two-hybrid (BACTH) Assay

The Bacterial Adenylate Cyclase Two-Hybrid (BACTH) system was employed as a genetic approach to assess protein-protein interactions. Gene inserts were amplified by PCR and cloned into pUT18C (CarbR) and pKT25 (KanR) vectors to generate N-terminal fusions and into pUT18 (CarbR) and pKNT25 (KanR) vectors to generate C-terminal fusions to the T18 and T25 fragments of adenylate cyclase, respectively. Miniprepped vectors (Qiagen plasmid miniprep kit) were then co-transformed into chemically competent *E. coli* BTH101 and outgrown with 500 μL LB for 1 hour at 37°C with shaking. Transformations were isolation streak plated on LB plates containing kanamycin 50 μg/mL and carbenicillin 100 μg/mL to select for transformants that had received both plasmids. Strains were then grown statically at 30°C from frozen stocks overnight in LB supplemented with kanamycin 50 μg/mL and carbenicillin 100 μg/mL and 3.5μL of each strain was then spotted onto an LB plate containing kanamycin 50 μg/mL, carbenicillin 100μg/mL, 0.5 mM IPTG and 40 μg/mL X-gal and incubated at 30°C for 48 hours prior to imaging on a HP Scanjet G4010 flatbed scanner.

### Western Blotting

Strains were grown to late log phase as described above in natural transformation assays +/- 0.2% arabinose where indicated. Cells were resuspended to an OD_600_ = 5 in IO + 100 μM IPTG and 0.2% arabinose where indicated and incubated for 1 hour statically. Cells were then concentrated to an OD_600_ = 50 in IO and mixed 1:1 with 2X SDS sample buffer [220 mM Tris pH 6.8, 25% glycerol, 1.8% SDS, 0.02% Bromophenol Blue, 5% β-mercaptoethanol] and boiled for 10 minutes at 100°C. Next, 10 μL of each sample was separated on a 10% SDS PAGE gel. Proteins were transferred to a PVDF membrane and incubated with mouse α- FLAG M2 monoclonal (Sigma) primary antibody. Then, blots were washed and incubated with a α-mouse secondary antibody conjugated to horseradish peroxidase (HRP) and developed using Pierce ECL Western blotting substrate. Blots were then imaged on a ProteinSimple FluorChem E instrument.

### Pilus fiber stretching assay

For purification of pili, a *V. cholerae* strains lacking the ability to make exopolysaccharides and all external appendages other than competence pili (i.e. mutants lacking flagella, MSHA pili, TCP pili and Vibrio polysaccharide (VPS) production) was grown to late log and labelled with AF488-mal dye exactly as described above, with the exception that cells were resuspended after washing to ∼10^10^ CFU/mL in 25 mM Tris pH 8.8. Cells were then loaded onto a SpinX 0.2 μm filter column (Sigma) and centrifuged at 16,000 x g for 1 minute and the flow through containing pili was collected. The flow through was centrifuged at 10,000 x g for 10 min at 4°C to remove cellular debris.

The magnetic tweezer setup was prepared as described previously (58). Briefly, cells were mounted on an Olympus IX81 inverted scope. Purified pilus filaments were incubated overnight with 1 μm MyOne carboxylated magnetic beads (Invitrogen) at a concentration allowing on average a single pilus to bind per bead. The beads were then allowed to settle onto a bed of hydrogel micropillars prepared as presented elsewhere (59). The surface was then scanned for the following configuration: a bead, with one pilus attached to it, stuck to one pillar with the pilus fiber attached to a neighboring pillar as well (**Fig. 5A**). About 10pN of pulling force was applied to the magnetic bead by using the magnetic tweezers. The images were prepared and analyzed using Fiji (57) and the lengths were calculated manually.

### Statistics

Statistical differences were assessed by Student’s t-test or one-way ANOVA tests followed by either a Dunnett’s or Tukey’s multiple comparisons post-test as indicated using GraphPad Prism software v. 9.0.2. For all transformation frequency and frequency of pilus retraction experiments statistical analyses were performed on log-transformed data. All statistical comparisons can be found in **Table S3**.

## Supplemental Information for

### This document includes

Figs. S1-S2

Movies S1-S8

Tables S1-S3

**Figure S1.**
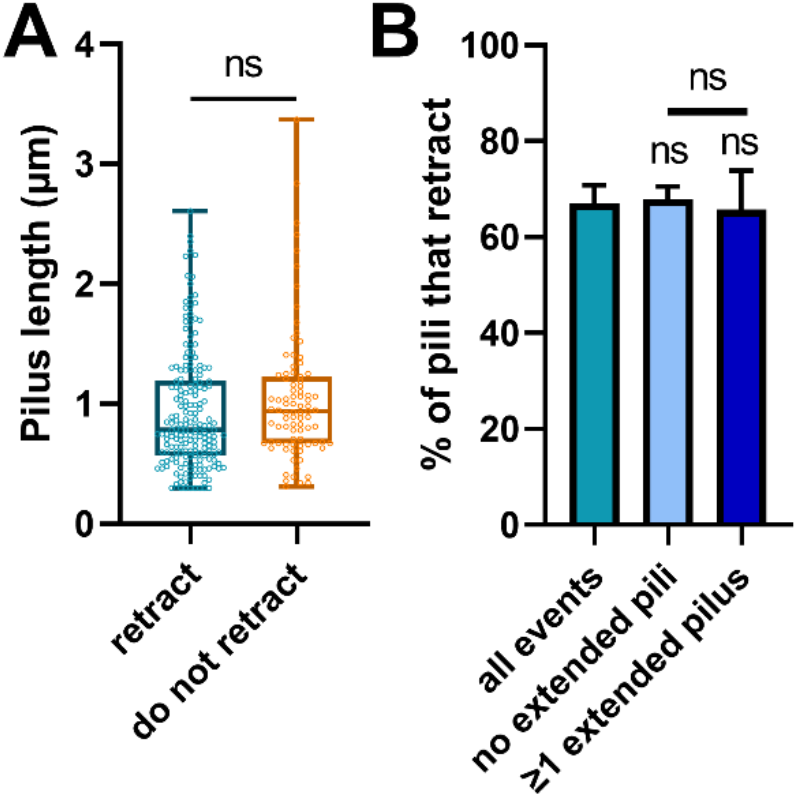
Neither pilus length nor presence of statically extended pili in cells affects likelihood that a pilus will retract. **(A)** pilus length measured after extension events seen in **Fig. 1E**. Data are from three independent biological replicates; *n* = 263. **(B)** The percent of pili that retract after extension was measured in cells that either displayed no extended pili or already had ≥1 statically extended pilus. The ‘all events’ bar is the same data as the ‘retract’ category in **Fig. 1E** and is included here for ease of comparison. Data were calculated from the same three independent biological replicates in **Fig. 1E**; ‘no extended pili’, *n* = 120. ‘≥1 extended pilus’, *n* = 143. Box plots represent the median and the upper and lower quartile, while the whiskers demarcate the range. Bar graphs are shown as the mean ± SD. Annotations directly above bars denote comparisons to the first bar in the graph. The comparison in **A** was made by Student’s t-test. Comparisons in **B** were made by one-way ANOVA with Tukey’s post test. ns, not significant.

**Figure S2.**
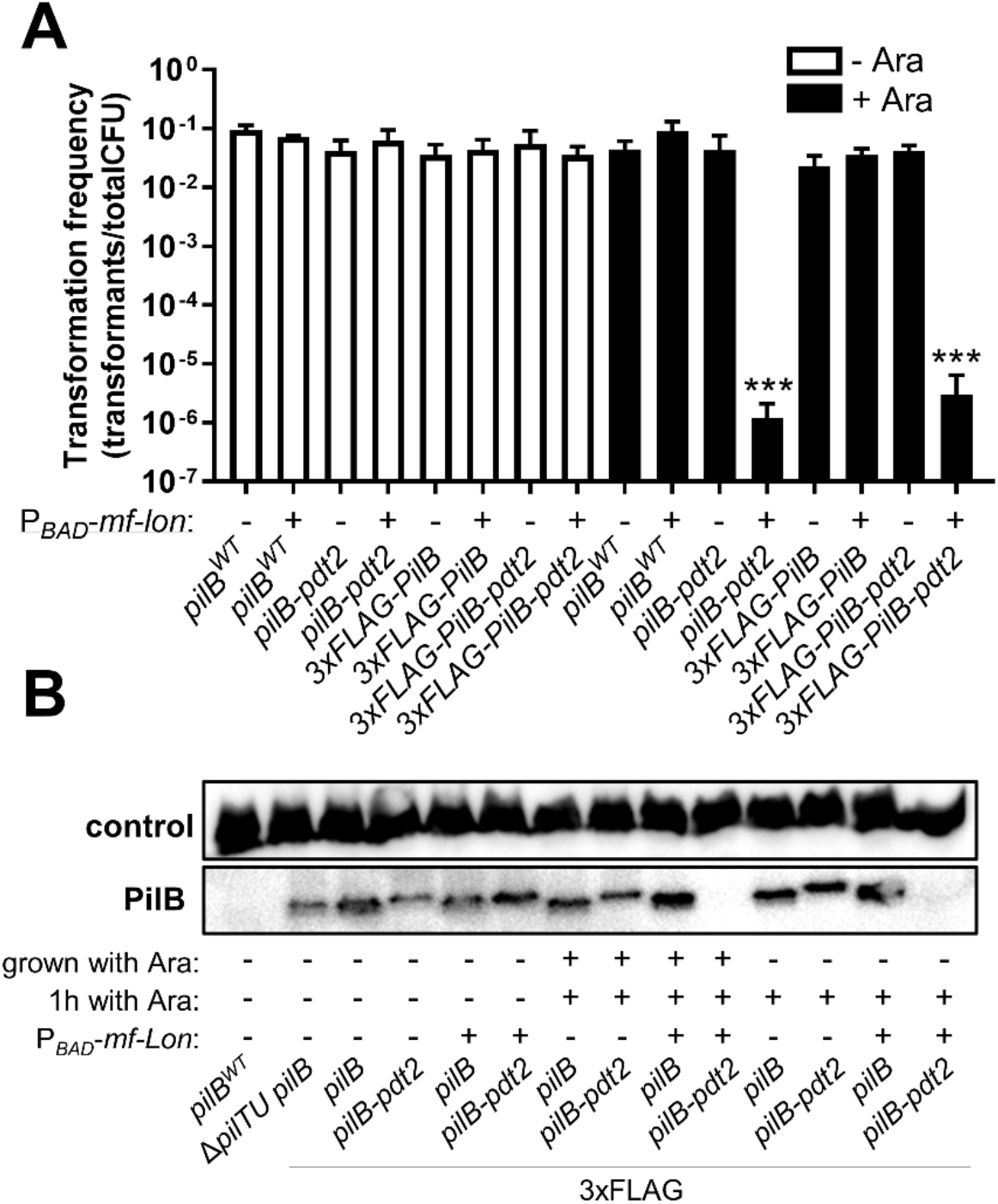
PilB with indicated translational fusions is functional and can be specifically degraded by mf-Lon protease. **(A)** Natural transformation assays of the indicated strains grown continually with or without 0.2% arabinose before addition of 200 ng tDNA. All strains, *n* = 3. **(B)** Western blot of the indicated strains to detect 3xFLAG tagged-PilB. Strains were grown continually with (grown with Ara; ‘+’) or without (grown with Ara; ‘-’) 0.2% arabinose and then incubated with (1h with Ara; ‘+’) or without (1h with Ara; ‘-’) 0.2% arabinose for 1 h as indicated. All strains except for *pilB*^*WT*^ contain an N-terminal 3xFLAG tag on PilB and strains that contain the P_*BAD*_*-mf-lon* ectopic expression construct are indicated (P_*BAD*_*-mf-lon*; ‘+’). Detection of a cross-reactive outer membrane porin that binds to the α-FLAG antibody (60) is shown as a loading control. Data are representative of two independent experiments. Bar graphs show the mean ± SD. Asterisks directly above bars denote comparisons to the comparable parent strain condition. All comparisons were made by one-way ANOVA with Dunnett’s post test. Ara, arabinose; *** = *P* < 0.001.

**Movie S1. Representative time-lapse of a Δ*pilTU* retraction event which was immediately preceded by extension**. Time-lapse of the montage shown in **Fig. 1A**, ‘yes’. The capture interval is 10 seconds between frames. Scale bar, 1 µm.

**Movie S2. Representative time-lapse of a Δ*pilTU* retraction event of an initially statically extended pilus**. Time-lapse of the montage shown in **Fig. 1A**, ‘no’. The capture interval is 10 seconds between frames. Scale bar, 1 µm.

**Movie S3. Representative time-lapse of a Δ*pilTU* Δ*pilB* P**_***BAD***_**-*pilB* strain grown without arabinose and incubated without arabinose**. Time-lapse of the montage shown in **Fig. 1C**, ‘no Ara’. The capture interval is 10 seconds between frames. Scale bar, 1 µm.

**Movie S4. Representative time-lapse of a Δ*pilTU* Δ*pilB* P**_***BAD***_**-*pilB* strain grown without arabinose and incubated with 0.2% arabinose for 30 min**. Time-lapse of the montage shown in **Fig. 1C**, ‘+ Ara 30 min’. The capture interval is 10 seconds between frames. Scale bar, 1 µm.

**Movie S5. Representative time-lapse of a Δ*pilTU* Δ*pilB* P**_***BAD***_**-*pilB* strain grown without arabinose and incubated with 0.2% arabinose for 3 h**. Time-lapse of the montage shown in **Fig. 1C**, ‘+ Ara 3 h’. The capture interval is 10 seconds between frames. Scale bar, 1 µm.

**Movie S6. Representative time-lapse of a Δ*pilTU* P**_***tac***_**-riboswitch-*pilT* strain grown without TP/IPTG and incubated with TP/IPTG during imaging**. Time-lapse of the montage shown in **Fig. 3B**, top panel. The capture interval is 10 seconds between frames. Scale bar, 1 µm

**Movie S7. Representative time-lapse of a Δ*pilTU* P**_***tac***_**-riboswitch-*pilT pilB***^***WT***^ **P**_***BAD***_***-mf*-*lon* strain grown without inducers, incubated with Ara for 1 h prior to imaging, and incubated with TP/IPTG/Ara during imaging**. Time-lapse of the montage shown in **Fig. 3B**, middle panel. The capture interval is 10 seconds between frames. Scale bar, 1 µm

**Movie S8. Representative time-lapse of a Δ*pilTU* P**_***tac***_**-riboswitch-*pilT pilB-*pdt2 P**_**BAD**_***-mf*-*lon* strain grown without inducers, incubated with Ara for 1 h prior to imaging, and incubated with TP/IPTG/Ara during imaging**. Time-lapse of the montage shown in **Fig. 3B**, bottom panel. The capture interval is 10 seconds between frames. Scale bar, 1 µm

**Table S1.**
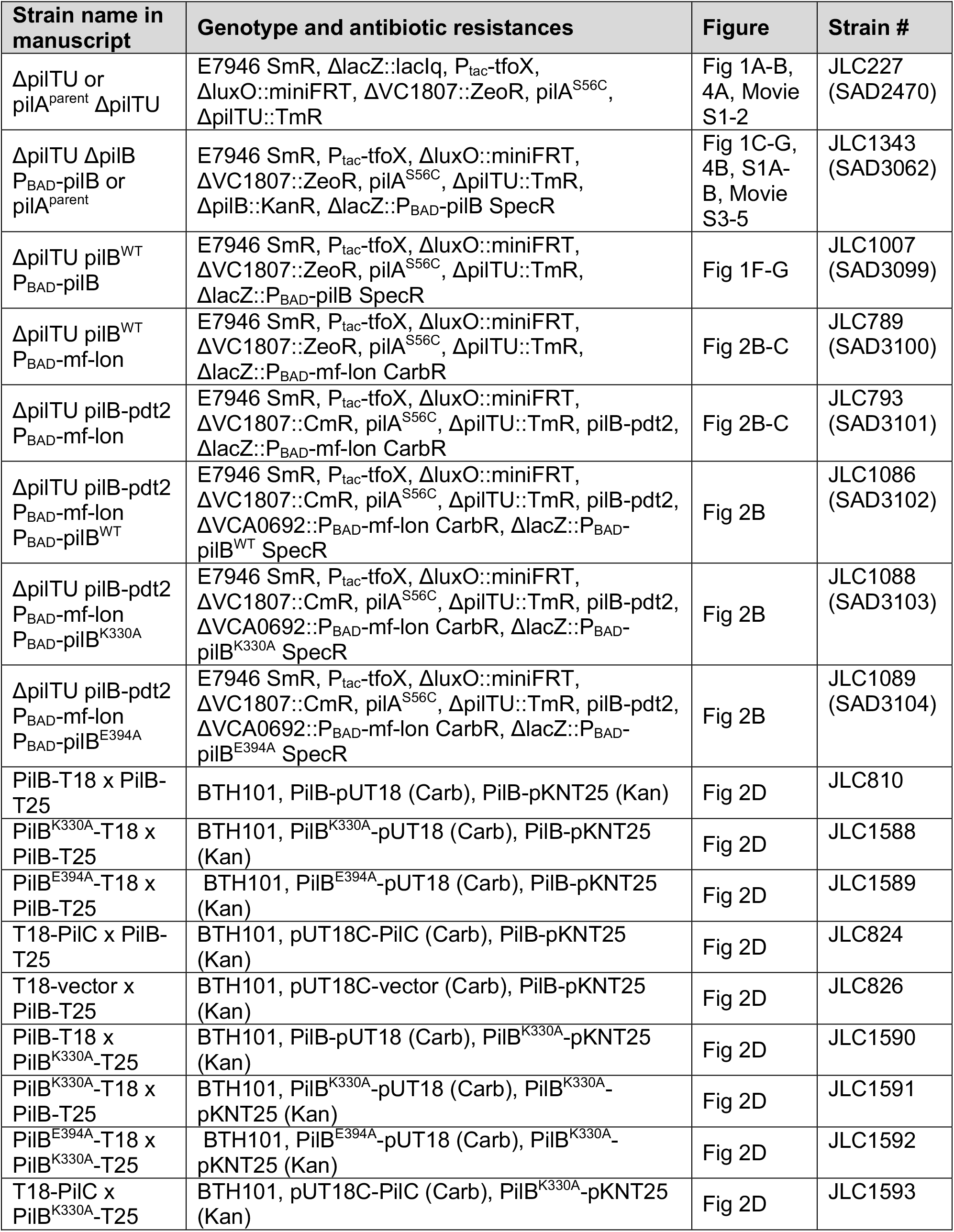
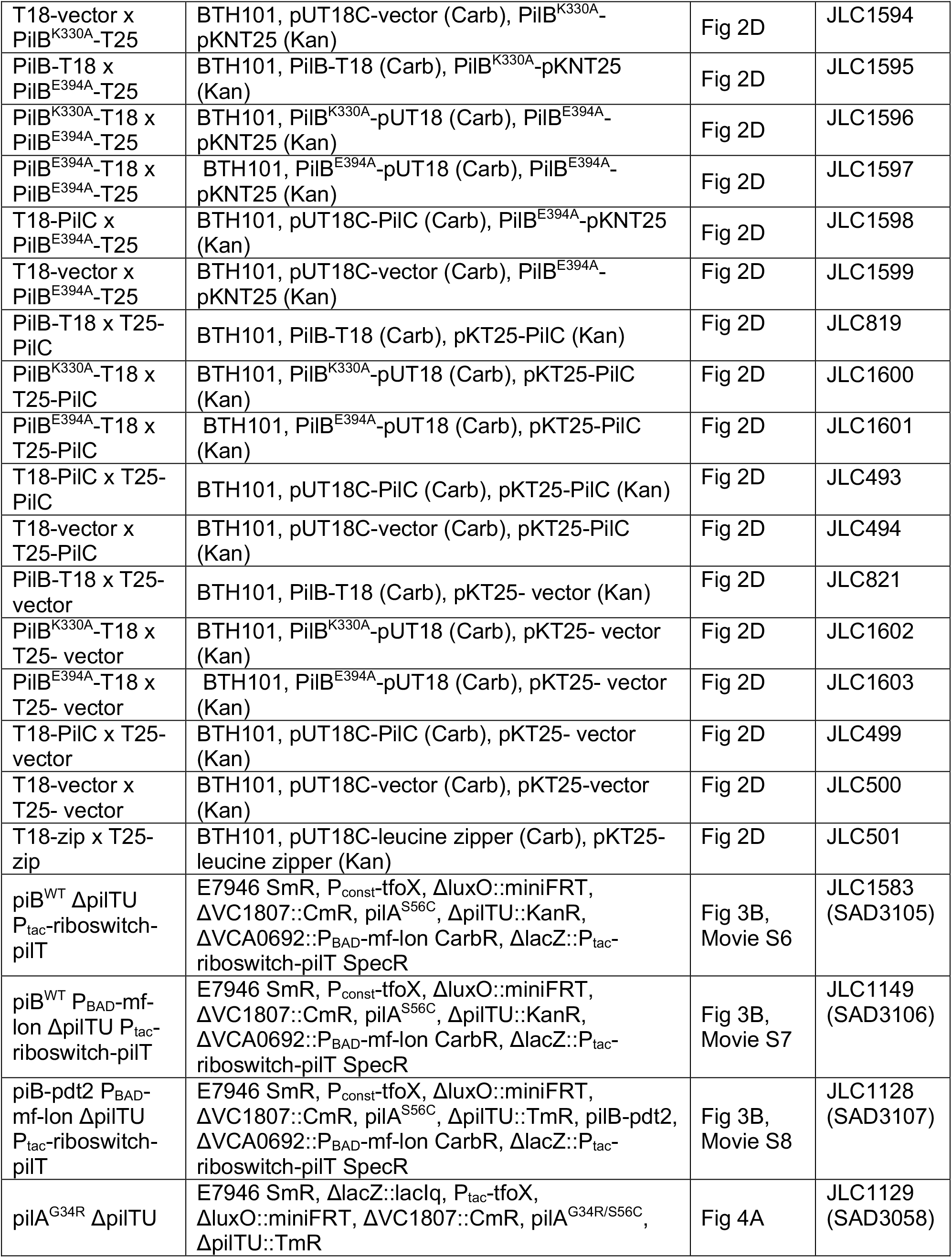
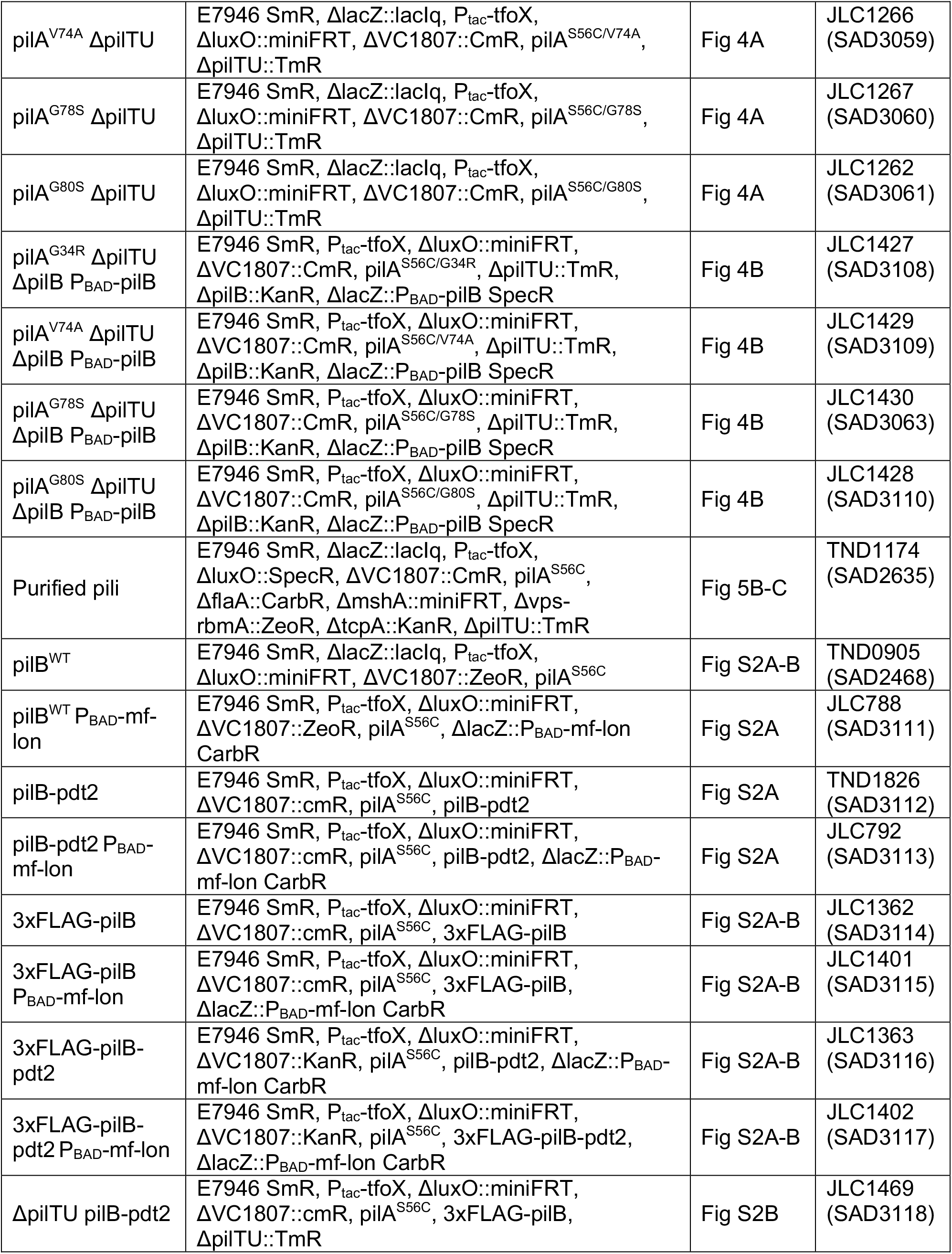
Strains used in this study.

**Table S2.**
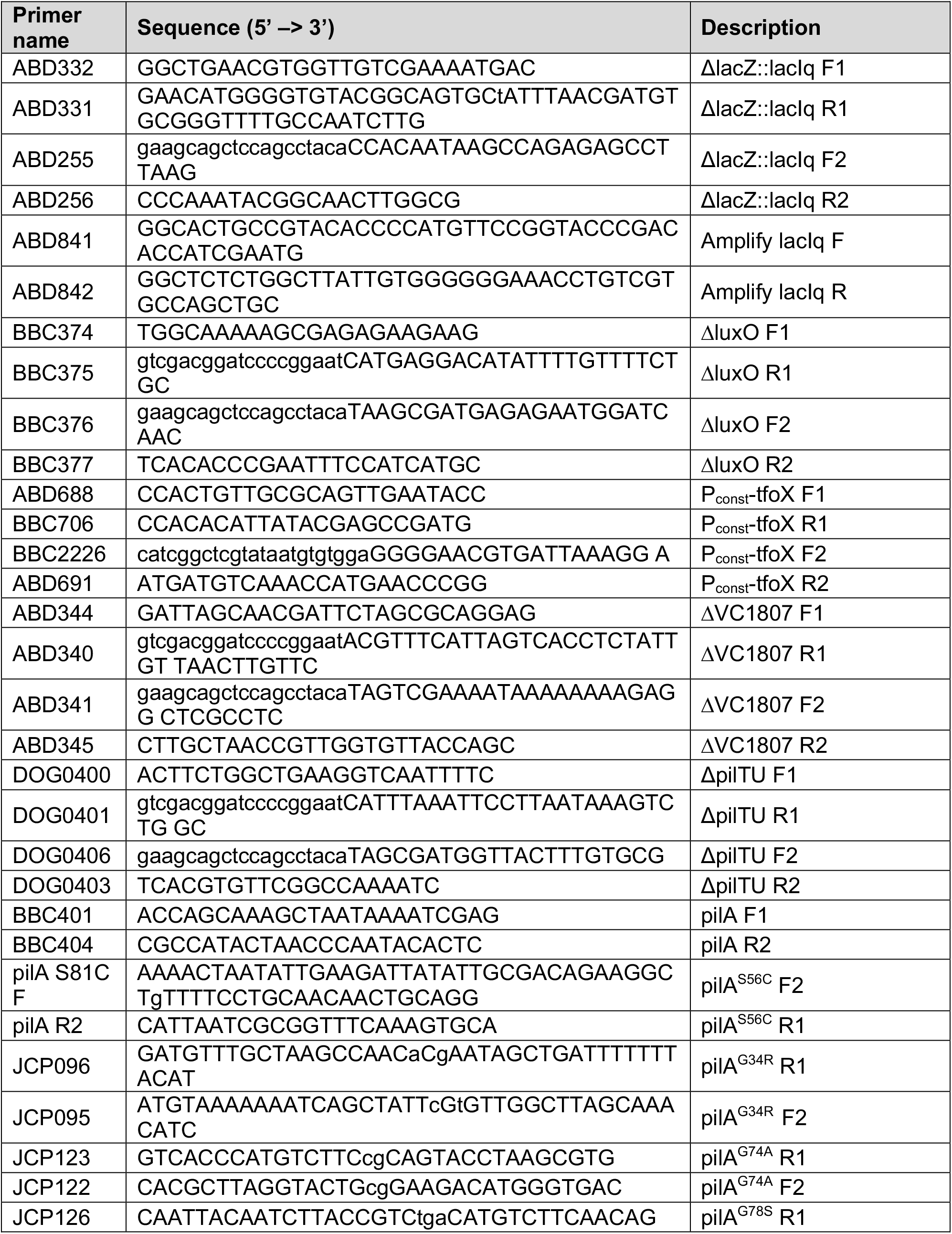
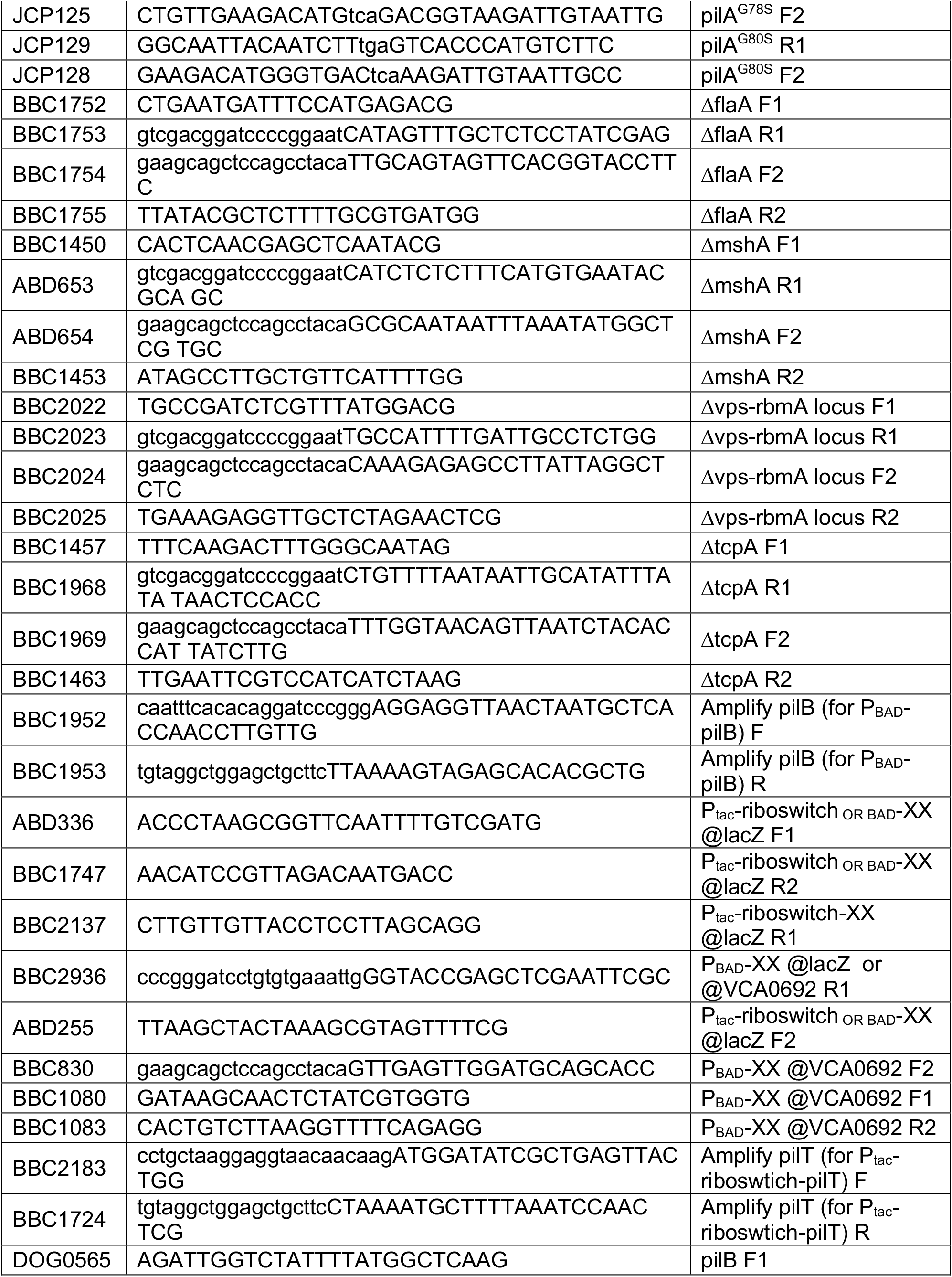
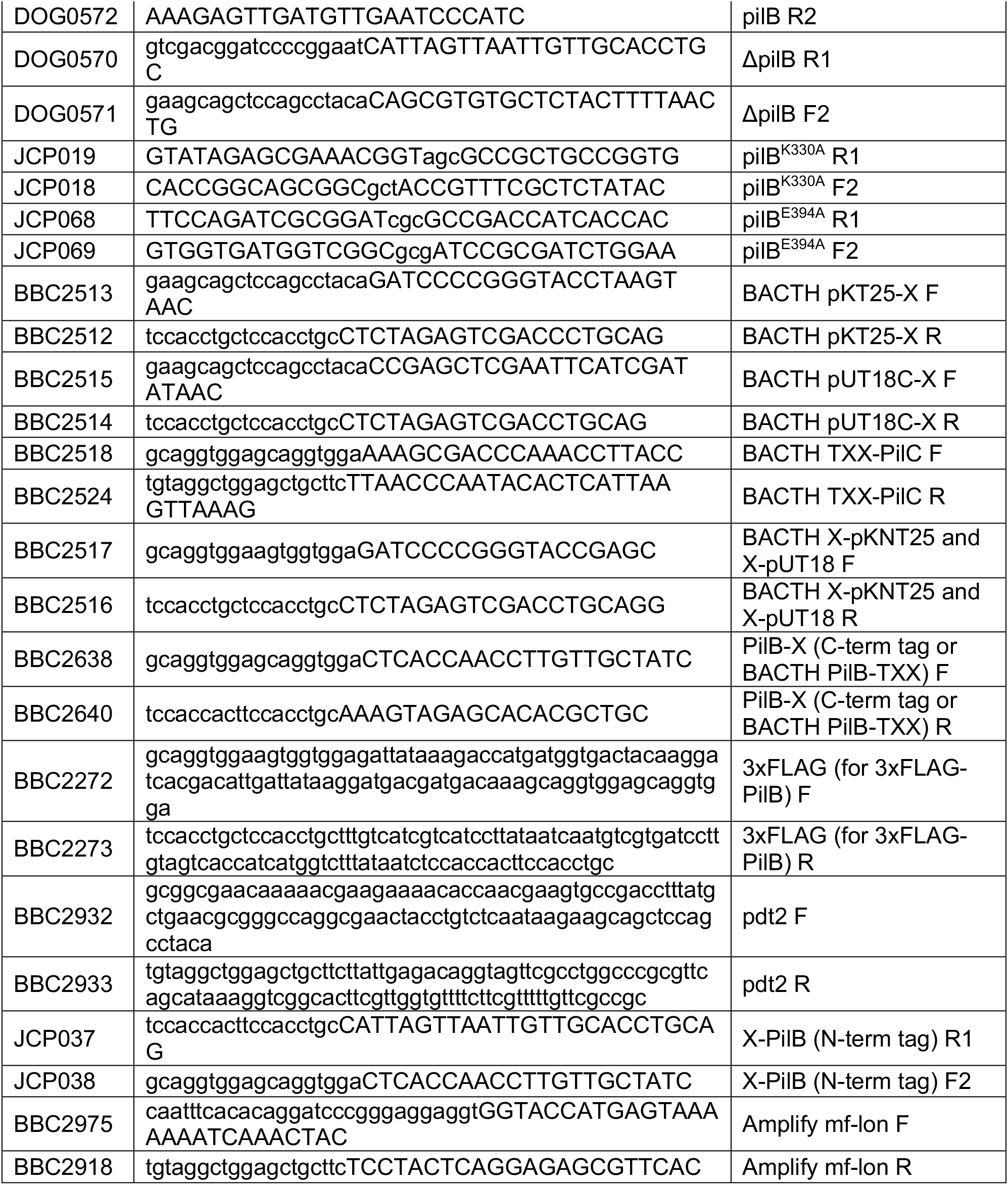
Primers used in this study.

**Table S3.** Statistical Comparisons.

| Comparison | P value<sup>#</sup> |  | Information |
| --- | --- | --- | --- |
| Figure 1B - Unpaired Two-tailed Student's T-test |  |  |  |
| ΔpilTU – Yes vs. No | **** | <0.0001 | Alpha = 0.05 |
| Figure 1D - Unpaired Two-tailed Student's T-test |  |  |  |
| ΔpilTU ΔpilB P<sub>BAD</sub>-pilB – 30min vs. 3hours | *** | 0.0002 | Alpha = 0.05 |
| Figure 1E - Unpaired Two-tailed Student's T-test |  |  |  |
| ΔpilTU ΔpilB P<sub>BAD</sub>-pilB – retract vs. do not retract | *** | 0.0003 | Alpha = 0.05 |
| Figure 1F - One way ANOVA with Tukey's multiple comparison test |  |  |  |
| pilB<sup>WT</sup> (-Ara, 0h) vs. ΔpilB (-Ara, 0h) | **** | <0.0001 | # of families = 1 |
| pilB<sup>WT</sup> (-Ara, 0h) vs. pilB<sup>WT</sup> (+Ara, 0h) | ns | 0.9594 | # of comparisons per family = 15 |
| pilB<sup>WT</sup> (-Ara, 0h) vs. ΔpilB (+Ara, 0h) | *** | 0.0002 | Alpha = 0.05 |
| pilB<sup>WT</sup> (-Ara, 0h) vs. pilB<sup>WT</sup> (+Ara, 3h) | ns | >0.9999 |  |
| pilB<sup>WT</sup> (-Ara, 0h) vs. ΔpilB (+Ara, 3h) | ns | >0.9999 |  |
| ΔpilB (-Ara, 0h) vs. pilB<sup>WT</sup> (+Ara, 0h) | **** | <0.0001 |  |
| ΔpilB (-Ara, 0h) vs. ΔpilB (+Ara, 0h) | **** | <0.0001 |  |
| ΔpilB (-Ara, 0h) vs. pilB<sup>WT</sup> (+Ara, 3h) | **** | <0.0001 |  |
| ΔpilB (-Ara, 0h) vs. ΔpilB (+Ara, 3h) | **** | <0.0001 |  |
| pilB<sup>WT</sup> (+Ara, 0h) vs. ΔpilB (+Ara, 0h) | **** | <0.0001 |  |
| pilB<sup>WT</sup> (+Ara, 0h) vs. pilB<sup>WT</sup> (+Ara, 3h) | ns | 0.9261 |  |
| pilB<sup>WT</sup> (+Ara, 0h) vs. ΔpilB (+Ara, 3h) | ns | 0.9772 |  |
| ΔpilB (+Ara, 0h) vs. pilB<sup>WT</sup> (+Ara, 3h) | *** | 0.0007 |  |
| ΔpilB (+Ara, 0h) vs. ΔpilB (+Ara, 3h) | *** | 0.0004 |  |
| pilB<sup>WT</sup> (+Ara, 3h) vs. ΔpilB (+Ara, 3h) | ns | >0.9999 |  |
| Figure 2B - One way ANOVA with Dunnett's multiple comparison test |  |  |  |
| (-Ara) pilB<sup>WT</sup> vs. pilB-pdt2 | ** | 0.0087 | # of families = 2 |
| (-Ara) pilB<sup>WT</sup> vs. pilB-pdt2 P<sub>BAD</sub>-pilB | ns | 0.7642 | # of comparisons per family = 4 |
| (-Ara) pilB<sup>WT</sup> vs. pilB-pdt2 P<sub>BAD</sub>-pilB<sup>K330A</sup> | * | 0.0128 | Alpha = 0.05 |
| (-Ara) pilB<sup>WT</sup> vs. pilB-pdt2 P<sub>BAD</sub>-pilB<sup>E394A</sup> | * | 0.0140 |  |
| (+Ara) pilB<sup>WT</sup> vs. pilB-pdt2 | ns | 0.8271 |  |
| (+Ara) pilB<sup>WT</sup> vs. pilB-pdt2 P<sub>BAD</sub>-pilB | ns | 0.9280 |  |
| (+Ara) pilB<sup>WT</sup> vs. pilB-pdt2 P<sub>BAD</sub>-pilB<sup>K330A</sup> | ns | 0.3919 |  |
| (+Ara) pilB<sup>WT</sup> vs. pilB-pdt2 P<sub>BAD</sub>-pilB<sup>E394A</sup> | ns | 0.9999 |  |
| Figure 4A - One way ANOVA with Dunnett's multiple comparison test |  |  |  |
| (Yes) pilA<sup>parent</sup> vs. pilA<sup>G34R</sup> | *** | 0.0002 | # of families = 2 |
| (Yes) pilA<sup>parent</sup> vs. pilA<sup>G80S</sup> | *** | 0.0008 | # of comparisons per family = 10 |
| (Yes) pilA<sup>parent</sup> vs. pilA<sup>V74A</sup> | * | 0.0208 | Alpha = 0.05 |
| (Yes) pilA<sup>parent</sup> vs. pilA<sup>G78S</sup> | ** | 0.0044 |  |
| (Yes) pilA<sup>G34R</sup> vs. pilA<sup>G80S</sup> | ns | 0.8918 |  |
| (Yes) pilA<sup>G34R</sup> vs. pilA<sup>V74A</sup> | ns | 0.0936 |  |
| (Yes) pilA<sup>G34R</sup> vs. pilA<sup>G78S</sup> | ns | 0.3558 |  |
| (Yes) pilA<sup>G80S</sup> vs. pilA<sup>V74A</sup> | ns | 0.3498 |  |
| (Yes) pilA<sup>G80S</sup> vs. pilA<sup>G78S</sup> | ns | 0.8339 |  |
| (Yes) pilA<sup>V74A</sup> vs. pilA<sup>G78S</sup> | ns | 0.8867 |  |
| (No) pilA<sup>parent</sup> vs. pilA<sup>G34R</sup> | *** | 0.0002 |  |
| (No) pilA<sup>parent</sup> vs. pilA<sup>G80S</sup> | *** | 0.0008 |  |
| (No) pilA<sup>parent</sup> vs. pilA<sup>V74A</sup> | * | 0.0208 |  |
| (No) pilA<sup>parent</sup> vs. pilA<sup>G78S</sup> | ** | 0.0044 |  |
| (No) pilA<sup>G34R</sup> vs. pilA<sup>G80S</sup> | ns | 0.8918 |  |
| (No) pilA<sup>G34R</sup> vs. pilA<sup>V74A</sup> | ns | 0.0936 |  |
| (No) pilA<sup>G34R</sup> vs. pilA<sup>G78S</sup> | ns | 0.3558 |  |
| (No) pilA<sup>G80S</sup> vs. pilA<sup>V74A</sup> | ns | 0.3498 |  |
| (No) pilA<sup>G80S</sup> vs. pilA<sup>G78S</sup> | ns | 0.8339 |  |
| (No) pilA<sup>V74A</sup> vs. pilA<sup>G78S</sup> | ns | 0.8867 |  |
| **Figure 4B - One way ANOVA with Tukey's multiple comparison test** |  |  |  |
| pilA<sup>parent</sup> (0h) vs. pilA<sup>G34R</sup> (0h) | ** | 0.0017 | # of families = 1 |
| pilA<sup>parent</sup> (0h) vs. pilA<sup>G80S</sup> (0h) | * | 0.0327 | # of comparisons per family = 45 |
| pilA<sup>parent</sup> (0h) vs. pilA<sup>V74A</sup> (0h) | *** | 0.0004 | Alpha = 0.05 |
| pilA<sup>parent</sup> (0h) vs. pilA<sup>G78S</sup> (0h) | ** | 0.0022 |  |
| pilA<sup>parent</sup> (0h) vs. pilA<sup>parent</sup> (3h) | **** | <0.0001 |  |
| pilA<sup>parent</sup> (0h) vs. pilA<sup>G34R</sup> (3h) | **** | <0.0001 |  |
| pilA<sup>parent</sup> (0h) vs. pilA<sup>G80S</sup> (3h) | ** | 0.0031 |  |
| pilA<sup>parent</sup> (0h) vs. pilA<sup>V74A</sup> (3h) | **** | <0.0001 |  |
| pilA<sup>parent</sup> (0h) vs. pilA<sup>G78S</sup> (3h) | ** | 0.0058 |  |
| pilA<sup>G34R</sup> (0h) vs. pilA<sup>G80S</sup> (0h) | ns | 0.9233 |  |
| pilA<sup>G34R</sup> (0h) vs. pilA<sup>V74A</sup> (0h) | ns | 0.9995 |  |
| pilA<sup>G34R</sup> (0h) vs. pilA<sup>G78S</sup> (0h) | ns | >0.9999 |  |
| pilA<sup>G34R</sup> (0h) vs. pilA<sup>parent</sup> (3h) | **** | <0.0001 |  |
| pilA<sup>G34R</sup> (0h) vs. pilA<sup>G34R</sup> (3h) | ns | 0.9289 |  |
| pilA<sup>G34R</sup> (0h) vs. pilA<sup>G80S</sup> (3h) | ns | >0.9999 |  |
| pilA<sup>G34R</sup> (0h) vs. pilA<sup>V74A</sup> (3h) | ns | 0.8918 |  |
| pilA<sup>G34R</sup> (0h) vs. pilA<sup>G78S</sup> (3h) | ns | 0.9999 |  |
| pilA<sup>G80S</sup> (0h) vs. pilA<sup>V74A</sup> (0h) | ns | 0.5941 |  |
| pilA<sup>G80S</sup> (0h) vs. pilA<sup>G78S</sup> (0h) | ns | 0.9530 |  |
| pilA<sup>G80S</sup> (0h) vs. pilA<sup>parent</sup> (3h) | **** | <0.0001 |  |
| pilA<sup>G80S</sup> (0h) vs. pilA<sup>G34R</sup> (3h) | ns | 0.2310 |  |
| pilA<sup>G80S</sup> (0h) vs. pilA<sup>G80S</sup> (3h) | ns | 0.9802 |  |
| pilA<sup>G80S</sup> (0h) vs. pilA<sup>V74A</sup> (3h) | ns | 0.1913 |  |
| pilA<sup>G80S</sup> (0h) vs. pilA<sup>G78S</sup> (3h) | ns | 0.9976 |  |
| pilA<sup>V74A</sup> (0h) vs. pilA<sup>G78S</sup> (0h) | ns | 0.9983 |  |
| pilA<sup>V74A</sup> (0h) vs. pilA<sup>parent</sup> (3h) | **** | <0.0001 |  |
| pilA<sup>V74A</sup> (0h) vs. pilA<sup>G34R</sup> (3h) | ns | 0.9992 |  |
| pilA<sup>V74A</sup> (0h) vs. pilA<sup>G80S</sup> (3h) | ns | 0.9931 |  |
| pilA<sup>V74A</sup> (0h) vs. pilA<sup>V74A</sup> (3h) | ns | 0.9976 |  |
| pilA<sup>V74A</sup> (0h) vs. pilA<sup>G78S</sup> (3h) | ns | 0.9604 |  |
| pilA<sup>G78S</sup> (0h) vs. pilA<sup>parent</sup> (3h) | **** | <0.0001 |  |
| pilA<sup>G78S</sup> (0h) vs. pilA<sup>G34R</sup> (3h) | ns | 0.8910 |  |
| pilA<sup>G78S</sup> (0h) vs. pilA<sup>G80S</sup> (3h) | ns | >0.9999 |  |
| pilA<sup>G78S</sup> (0h) vs. pilA<sup>V74A</sup> (3h) | ns | 0.8444 |  |
| pilA<sup>G78S</sup> (0h) vs. pilA<sup>G78S</sup> (3h) | ns | >0.9999 |  |
| pilA<sup>parent</sup> (3h) vs. pilA<sup>G34R</sup> (3h) | **** | <0.0001 |  |
| pilA<sup>parent</sup> (3h) vs. pilA<sup>G80S</sup> (3h) | **** | <0.0001 |  |
| pilA<sup>parent</sup> (3h) vs. pilA<sup>V74A</sup> (3h) | **** | <0.0001 |  |
| pilA<sup>parent</sup> (3h) vs. pilA<sup>G78S</sup> (3h) | **** | <0.0001 |  |
| pilA<sup>G34R</sup> (3h) vs. pilA<sup>G80S</sup> (3h) | ns | 0.8200 |  |
| pilA<sup>G34R</sup> (3h) vs. pilA<sup>V74A</sup> (3h) | ns | >0.9999 |  |
| pilA<sup>G34R</sup> (3h) vs. pilA<sup>G78S</sup> (3h) | ns | 0.6612 |  |
| pilA<sup>G80S</sup> (3h) vs. pilA<sup>V74A</sup> (3h) | ns | 0.7616 |  |
| pilA<sup>G80S</sup> (3h) vs. pilA<sup>G78S</sup> (3h) | ns | >0.9999 |  |
| pilA<sup>V74A</sup> (3h) vs. pilA<sup>G78S</sup> (3h) | ns | 0.5927 |  |
| **Figure 5C – Unpaired Two-tailed Student's T-test** |  |  |  |
| magnet OFF vs. magnet ON | **** | <0.0001 | Alpha = 0.05 |
| **Supplemental Figure 1A – Unpaired Two-tailed Student's T-test** |  |  |  |
| (pilus length) retract vs. do not retract | ns | 0.1046 | Alpha = 0.05 |
| **Supplemental Figure 1B – One way ANOVA with Tukey's multiple comparison test** |  |  |  |
| all events vs. no extended pili | ns | 0.9806 | # of families = 1 |
| all events vs. ≥1 extended pilus | ns | 0.9494 | # of comparisons per family = 3 |
| no extended pili vs. ≥1 extended pilus | ns | 0.8755 | Alpha = 0.05 |
| **Supplemental Figure 2A - One way ANOVA with Dunnett's multiple comparison test** |  |  |  |
| (-Ara) pilB<sup>WT</sup> vs. pilB<sup>WT</sup> P<sub>BAD-mf-lon</sub> | ns | 0.9964 | # of families = 2 |
| (-Ara) pilB<sup>WT</sup> vs. pilB-pdt2 P<sub>BAD-mf-lon</sub> | ns | 0.8439 | # of comparisons per family = 28 |
| (-Ara) pilB<sup>WT</sup> vs. pilB-pdt2 | ns | 0.3288 | Alpha = 0.05 |
| (-Ara) pilB<sup>WT</sup> vs. 3xFLAG-PilB | ns | 0.1885 |  |
| (-Ara) pilB<sup>WT</sup> vs. 3xFLAG-PilB-pdt2 | ns | 0.6223 |  |
| (-Ara) pilB<sup>WT</sup> vs. 3xFLAG-PilB P<sub>BAD-mf-lon</sub> | ns | 0.3960 |  |
| (-Ara) pilB<sup>WT</sup> vs. 3xFLAG-PilB-pdt2 P<sub>BAD-mf-lon</sub> | ns | 0.1902 |  |
| (-Ara) pilB<sup>WT</sup> P<sub>BAD-mf-lon</sub> vs. pilB-pdt2 P<sub>BAD-mf-lon</sub> | ns | 0.9950 |  |
| (-Ara) pilB<sup>WT</sup> P<sub>BAD-mf-lon</sub> vs. pilB-pdt2 | ns | 0.7083 |  |
| (-Ara) pilB<sup>WT</sup> P<sub>BAD-mf-lon</sub> vs. 3xFLAG-PilB | ns | 0.4936 |  |
| (-Ara) pilB<sup>WT</sup> P<sub>BAD-mf-lon</sub> vs. 3xFLAG-PilB-pdt2 | ns | 0.9404 |  |
| (-Ara) pilB<sup>WT</sup> P<sub>BAD-mf-lon</sub> vs. 3xFLAG-PilB P<sub>BAD-mf-lon</sub> | ns | 0.7832 |  |
| (-Ara) pilB<sup>WT</sup> P<sub>BAD-mf-lon</sub> vs. 3xFLAG-PilB-pdt2 P<sub>BAD-mf-lon</sub> | ns | 0.4968 |  |
| (-Ara) pilB-pdt2 P<sub>BAD-mf-lon</sub> vs. pilB-pdt2 | ns | 0.9764 |  |
| (-Ara) pilB-pdt2 P<sub>BAD-mf-lon</sub> vs. 3xFLAG-PilB | ns | 0.8826 |  |
| (-Ara) pilB-pdt2 P<sub>BAD-mf-lon</sub> vs. 3xFLAG-PilB-pdt2 | ns | 0.9999 |  |
| (-Ara) pilB-pdt2 P<sub>BAD-mf-lon</sub> vs. 3xFLAG-PilB P<sub>BAD-mf-lon</sub> | ns | 0.9902 |  |
| (-Ara) *pilB*-pdt2 $P_{BAD}$-mf-lon vs. 3xFLAG-*PilB*-pdt2 $P_{BAD}$-mf-lon | ns | 0.8847 | |
| (-Ara) *pilB*-pdt2 vs. 3xFLAG-*PilB* | ns | >0.9999 |  |
| (-Ara) *pilB*-pdt2 vs. 3xFLAG-*PilB*-pdt2 | ns | 0.9992 |  |
| (-Ara) *pilB*-pdt2 vs. 3xFLAG-*PilB* $P_{BAD}$-mf-lon | ns | >0.9999 | |
| (-Ara) *pilB*-pdt2 vs. 3xFLAG-*PilB*-pdt2 $P_{BAD}$-mf-lon | ns | >0.9999 | |
| (-Ara) 3xFLAG-*PilB* vs. 3xFLAG-*PilB*-pdt2 | ns | 0.9817 |  |
| (-Ara) 3xFLAG-*PilB* vs. 3xFLAG-*PilB* $P_{BAD}$-mf-lon | ns | 0.9995 | |
| (-Ara) 3xFLAG-*PilB* vs. 3xFLAG-*PilB*-pdt2 $P_{BAD}$-mf-lon | ns | >0.9999 | |
| (-Ara) 3xFLAG-*PilB*-pdt2 vs. 3xFLAG-*PilB* $P_{BAD}$-mf-lon | ns | 0.9999 | |
| (-Ara) 3xFLAG-*PilB*-pdt2 vs. 3xFLAG-*PilB*-pdt2 $P_{BAD}$-mf-lon | ns | 0.9824 | |
| (-Ara) 3xFLAG-*PilB* $P_{BAD}$-mf-lon vs. 3xFLAG-*PilB*-pdt2 $P_{BAD}$-mf-lon | ns | 0.9995 | |
| (+Ara) *pilB*<sup>WT</sup> vs. *pilB*<sup>WT</sup> $P_{BAD}$-mf-lon | ns | 0.9728 | |
| (+Ara) *pilB*<sup>WT</sup> vs. *pilB*-pdt2 $P_{BAD}$-mf-lon | **** | <0.0001 | |
| (+Ara) *pilB*<sup>WT</sup> vs. *pilB*-pdt2 | ns | >0.9999 |  |
| (+Ara) *pilB*<sup>WT</sup> vs. 3xFLAG-*PilB* | ns | 0.9724 |  |
| (+Ara) *pilB*<sup>WT</sup> vs. 3xFLAG-*PilB*-pdt2 | ns | >0.9999 |  |
| (+Ara) *pilB*<sup>WT</sup> vs. 3xFLAG-*PilB* $P_{BAD}$-mf-lon | ns | >0.9999 | |
| (+Ara) *pilB*<sup>WT</sup> vs. 3xFLAG-*PilB*-pdt2 $P_{BAD}$-mf-lon | **** | <0.0001 | |
| (+Ara) *pilB*<sup>WT</sup> $P_{BAD}$-mf-lon vs. *pilB*-pdt2 $P_{BAD}$-mf-lon | **** | <0.0001 | |
| (+Ara) *pilB*<sup>WT</sup> $P_{BAD}$-mf-lon vs. *pilB*-pdt2 | ns | 0.8779 | |
| (+Ara) *pilB*<sup>WT</sup> $P_{BAD}$-mf-lon vs. 3xFLAG-*PilB* | ns | 0.5460 | |
| (+Ara) *pilB*<sup>WT</sup> $P_{BAD}$-mf-lon vs. 3xFLAG-*PilB*-pdt2 | ns | 0.9706 | |
| (+Ara) *pilB*<sup>WT</sup> $P_{BAD}$-mf-lon vs. 3xFLAG-*PilB* $P_{BAD}$-mf-lon | ns | 0.9121 | |
| (+Ara) *pilB*<sup>WT</sup> $P_{BAD}$-mf-lon vs. 3xFLAG-*PilB*-pdt2 $P_{BAD}$-mf-lon | **** | <0.0001 | |
| (+Ara) *pilB*-pdt2 $P_{BAD}$-mf-lon vs. *pilB*-pdt2 | **** | <0.0001 | |
| (+Ara) *pilB*-pdt2 $P_{BAD}$-mf-lon vs. 3xFLAG-*PilB* | **** | <0.0001 | |
| (+Ara) *pilB*-pdt2 $P_{BAD}$-mf-lon vs. 3xFLAG-*PilB*-pdt2 | **** | <0.0001 | |
| (+Ara) *pilB*-pdt2 $P_{BAD}$-mf-lon vs. 3xFLAG-*PilB* $P_{BAD}$-mf-lon | **** | <0.0001 | |
| (+Ara) *pilB*-pdt2 $P_{BAD}$-mf-lon vs. 3xFLAG-*PilB*-pdt2 $P_{BAD}$-mf-lon | ns | 0.9867 | |
| (+Ara) *pilB*-pdt2 vs. 3xFLAG-*PilB* | ns | 0.9980 |  |
| (+Ara) *pilB*-pdt2 vs. 3xFLAG-*PilB*-pdt2 | ns | >0.9999 |  |
| (+Ara) *pilB*-pdt2 vs. 3xFLAG-*PilB* $P_{BAD}$-mf-lon | ns | >0.9999 | |
| (+Ara) *pilB*-pdt2 vs. 3xFLAG-*PilB*-pdt2 $P_{BAD}$-mf-lon | **** | <0.0001 | |
| (+Ara) 3xFLAG-*PilB* vs. 3xFLAG-*PilB*-pdt2 | ns | 0.9746 |  |
| (+Ara) 3xFLAG-*PilB* vs. 3xFLAG-*PilB* $P_{BAD}$-mf-lon | ns | 0.9953 | |
| (+Ara) 3xFLAG-*PilB* vs. 3xFLAG-*PilB*-pdt2 $P_{BAD}$-mf-lon | **** | <0.0001 | |
| (+Ara) 3xFLAG-*PilB*-pdt2 vs. 3xFLAG-*PilB* $P_{BAD}$-mf-lon | ns | >0.9999 | |
| (+Ara) 3xFLAG-PilB-pdt2 vs. 3xFLAG-PilB-pdt2 $P_{BAD}$-mf-lon | **** | <0.0001 | |
| (+Ara) 3xFLAG-PilB $P_{BAD}$-mf-lon vs. 3xFLAG-PilB-pdt2 $P_{BAD}$-mf-lon | **** | <0.0001 | |

#Statistical differences were assessed using GraphPad Prism v.9.0.2 software. For all transformation frequency experiments and frequency of pilus retraction experiments, statistical analyses were performed on the log transformed data (i.e. **Figs 1D, 1F, 2B, 4B and S2A**).

## Notes

### Competing Interest Statement

The authors have declared no competing interest.

